# Mechanosensory trichome cells evoke a mechanical stimuli–induced immune response in plants

**DOI:** 10.1101/2021.06.13.448005

**Authors:** Mamoru Matsumura, Mika Nomoto, Tomotaka Itaya, Yuri Aratani, Mizuki Iwamoto, Takakazu Matsuura, Yuki Hayashi, Toshinori Kinoshita, Izumi C Mori, Takamasa Suzuki, Shigeyuki Betsuyaku, Steven H Spoel, Masatsugu Toyota, Yasuomi Tada

## Abstract

Perception of pathogen-derived ligands by corresponding host receptors is a pivotal strategy in eukaryotic innate immunity. In plants, this is complemented by circadian anticipation of infection timing, promoting basal resistance even in the absence of pathogen threat. Here, we report that trichomes, hair-like structures on the epidermis, directly sense external mechanical forces caused by raindrops to anticipate waterborne infections in *Arabidopsis thaliana*. Exposure of leaf surfaces to mechanical stimuli initiates the concentric propagation of intercellular calcium waves away from trichomes to induce defence-related genes. Propagating calcium waves enable effective immunity against pathogenic microbes through the calmodulin-binding transcription activator 3 (CAMTA3) and mitogen-activated protein kinases. We propose a novel layer of plant immunity in which trichomes function as mechanosensory cells to detect potential risks.

## Introduction

Innate immunity is an evolutionarily conserved front line of defence across the plant and animal kingdoms. In plants, pattern-recognition receptors (PRRs), such as leucine-rich repeat receptor-like kinases (LRR-RLKs) and LRR receptor proteins (LRR-RPs), specifically recognize microbe-associated molecular patterns (MAMPs) as non-self molecules, leading to the activation of pattern-triggered immunity (PTI) to limit pathogen proliferation^1, 2^. While adapted pathogens have evolved virulence effectors that can circumvent PTI, plants also deploy disease resistance (*R*) genes, primarily encoding nucleotide-binding LRR proteins, which mount effector-triggered immunity (ETI)^3–5^. ETI often culminates in a hypersensitive response as well as acute and localized cell death induced at the site of infection and accompanied by profound transcriptional changes of defence-related genes to retard pathogen growth^4, 5^. These ligand-receptor systems are largely dependent on a transient increase in intracellular calcium concentration ([Ca^2+^]_i_), followed by the initiation of phosphorylation-dependent signalling cascades, including mitogen-activated protein kinases (MAPKs) and calcium-dependent protein kinases, that orchestrate a complex transcriptional network and the activity of immune mediators^6, 7^.

In addition to PTI and ETI, plant immunity can be induced periodically in the absence of pathogen threat, a process that is under the control of the circadian clock and driven by daily oscillations in humidity as well as light-dark cycles^8–10^. Such responses enable plants to prepare for the potential increased risk of infection at the time when microbes are anticipated to be most infectious. Therefore, the anticipation of potentially pathogenic microorganisms through sensing of climatological changes on the one hand and their specific detection on the other constitute two distinct layers of the plant immune system.

Among the climatological factors that affect the outcome of plant-microbe interactions, rain is a major cause of devastating plant diseases, as fungal spores and bacteria are spread through rain-dispersed aerosols or ballistic particles splashed from neighbouring infected plants. In addition, raindrops contain plant pathogens, including *Pseudomonas syringae* and *Xanthomonas campestris*, and negatively regulate stomatal closure, which facilitates pathogen entry into leaf tissues^11–13^. High humidity, which is usually associated with rain, enhances the effects of bacterial pathogen effectors, such as HopM1, and establishes an aqueous apoplast for aggressive host colonization^14^. These findings suggest that it would be beneficial for plants to recognize rain as an early risk factor for infectious diseases.

How do plants respond to rain? Rain induces the expression of mechanosensitive *TOUCH* (*TCH*) genes in plants^15^. Mechanostimulation may affect a variety of plant physiological processes mediated by hormones such as auxin, ethylene, and gibberellin^16–19^. *Arabidopsis thaliana* seedlings exposed to rain-simulating water spray accumulate the immune phytohormone jasmonic acid (JA) to promote the expression of JA-responsive genes^20^. Thus, rain modulates both mechanotransduction and hormone-signalling pathways that could affect the growth and development of plants as well as environmental responses. However, the regulatory mechanisms underpinning the rain-activated signalling pathway have not been fully elucidated.

Here, we report a novel layer of the plant immune system evoked by sensing mechanostimulation of falling raindrops: trichomes, hair-like cells on the leaf surface, function as mechanosensory cells that mount an effective immune response against both biotrophic and necrotrophic pathogens. When trichomes are mechanically stimulated, intercellular calcium waves are concentrically propagated away from them, and this is followed by the activation of MAPKs and initiation of the calcium- and calmodulin-binding transcription activator (CAMTA)-regulated immune response. We propose that plants directly recognize rain as a risk factor and evoke a rapid immune response that substantially contributes to early detection of, and protection from, potential pathogens.

## Results and discussion

### Rain and mechanical stimuli induce mechanosensitive genes involved in plant immunity

To investigate the effect of rain on transcriptional changes in *Arabidopsis* leaves, we performed transcriptome deep sequencing (RNA-seq) of wild-type Columbia (Col-0) *Arabidopsis* treated with artificial raindrops (Supplementary Fig. 1a; Methods). After applying only 10 falling droplets, we detected the marked induction of 1,050 genes 15 min after treatment (Supplementary Table 1). Gene Ontology (GO) analysis of these genes revealed a striking enrichment in categories associated with plant immunity, as evidenced by the expression of major immune regulators including *WRKY DNA-BINDING PROTEIN* (*WRKY*) genes, *CALMODULIN BINDING PROTEIN 60-LIKE* g (*CBP60g*), *MYB DOMAIN PROTEIN* (*MYB*) genes, *ETHYLENE RESPONSE FACTOR* (*ERF*) genes, and *MAP KINASE* (*MPK*) genes^21, 22^ (Fig. 1a, Supplementary Table 1, Supplementary Table 2). The touch-induced genes *TCH2* and *TCH4* were also highly upregulated in response to one falling raindrop (falling) compared to a water droplet placed directly on the leaf surface (static) (Supplementary Fig. 1b). These results suggested that mechanosensation is involved in altering transcriptional activity.

**Fig. 1.**
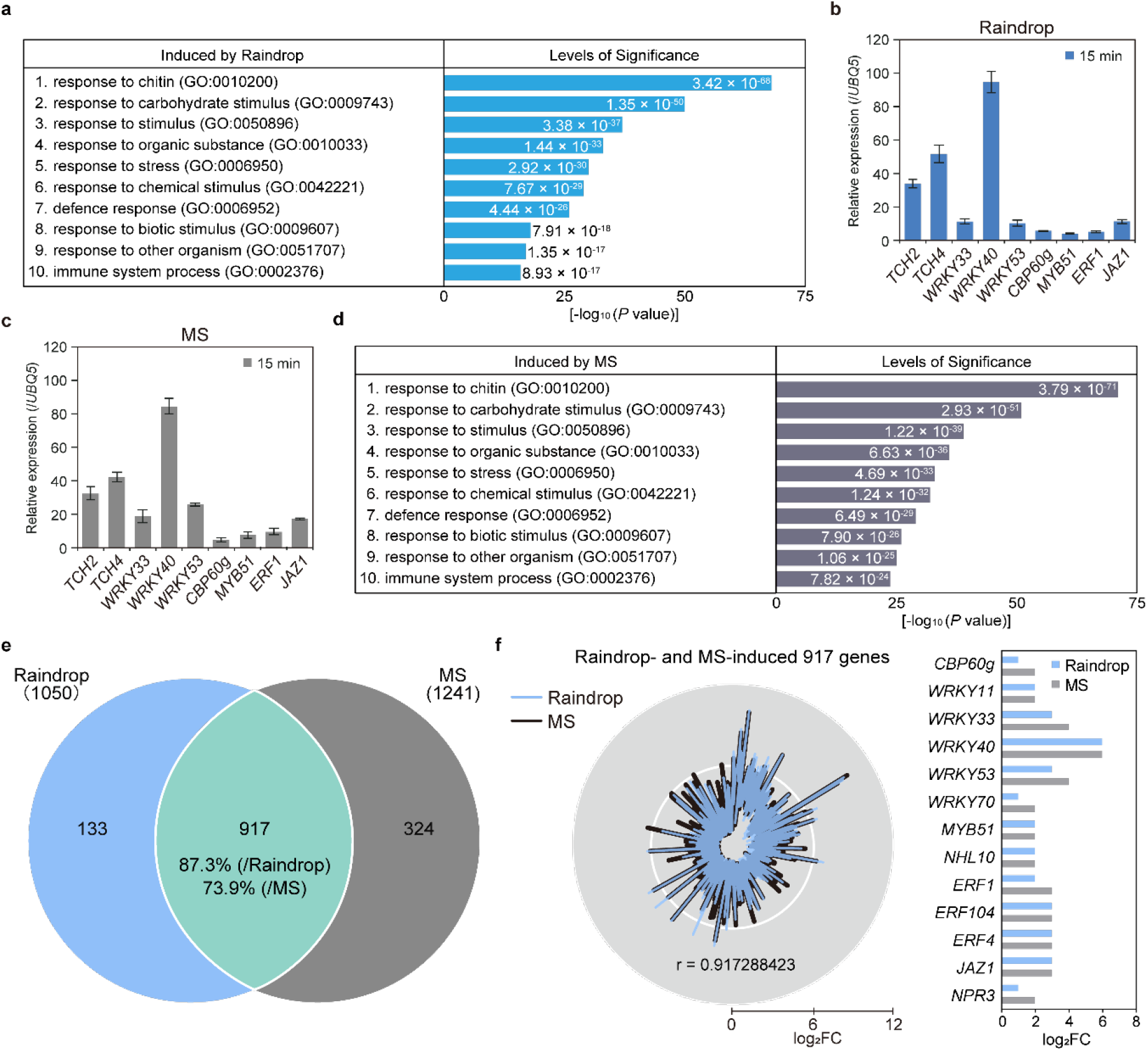
Raindrop-induced gene expression highly correlates with mechanical stimuli (MS)-induced gene expression in *Arabidopsis*. **a** Enriched Gene Ontology (GO) categories of 1,050 raindrop (10 droplets)-induced genes in the wild type (Col-0). The top 10 categories are shown in ascending order of *P* values. **b**, **c** Transcript levels of MS-induced and defence-related genes in 4-week-old Col-0 plants 15 min after being treated with 10 falling droplets (raindrop, **b**) or 1 brushing (MS, **c**), determined by RT-qPCR and normalized to *UBIQUITIN 5* (*UBQ5*). Data are presented as mean ± SD. **d** Enriched GO categories of 1,241 MS (1 brushing)-induced genes in Col-0. The top 10 categories are shown in ascending order of *P* values. **e** Venn diagram of the overlap between transcriptome datasets from raindrop- and MS-induced genes (*P* < 0.05). **f** Radar chart of intensity compared with mock (log_2_FC) and Pearson correlation coefficient (r = 0.917288423) of 917 raindrop- and MS-induced genes (left). Intensities of major immune regulator genes induced by raindrops and MS in RNA-seq analysis (log_2_FC) (right).

To validate this hypothesis, we mechanically stimulated rosette leaves by gently brushing them one to ten times along the main veins with a small paint brush (Supplementary Fig. 1c; Methods) and analysed the expression profile of the immune regulator *WRKY33*, which was responsive to raindrops. *WRKY33* expression was maximally induced 15 min after brushing the leaves one to four times (Supplementary Fig. 1d). Next, we compared gene expression patterns between leaves that were brushed once and those that received 10 falling raindrops. Both raindrops and brushing strongly upregulated *TCH2*, *TCH4*, *WRKY33*, *WRKY40*, *WRKY53*, *CBP60g*, *MYB51*, *ERF1*, and *JASMONATE-ZIM-DOMAIN PROTEIN 1* (*JAZ1*) expression ^15, 21, 22^ (Fig. 1b, c), suggesting that raindrops are likely recognized as a mechanical stimulus.

Then, to comprehensively identify mechanosensitive genes, we performed an RNA-seq analysis of leaves brushed once. We identified 1,241 genes that were significantly induced 15 min after this treatment relative to control plants (Supplementary Table 3). These mechanical stimuli (MS)-induced genes were primarily categorized as plant immune responses, such as response to chitin, defence response, and immune system response (Fig. 1d, Supplementary Table 4). We found that 87.3% of raindrop-induced genes and 73.9% of MS-induced genes overlapped (Fig. 1e): this set of 917 genes expressed upon both treatments was enriched in GO categories associated with stress responses (Supplementary Fig. 1e). Furthermore, the expression levels of these 917 genes, including major immune regulators, were strongly positively correlated between the two treatments (Pearson correlation coefficient r = 0.917) (Fig. 1f). These transcriptome analyses indicated that falling raindrops stimulate the expression of mechanosensitive genes involved in environmental stress responses, including plant immunity.

### Rain and MS rapidly activate plant immune responses

To further characterize raindrop-induced genes, we conducted a comparative analysis with published transcriptome datasets. Many raindrop- and MS-induced genes were also expressed during major plant immune responses, such as those triggered by the immune phytohormones salicylic acid (SA), which is effective against biotrophic pathogens (21%; 193/917 genes), and JA, which mounts immune responses to necrotrophic pathogens (11.8%; 108/917 genes); the bacterial-derived peptide flg22, which activates PTI (37%; 339/917 genes); and the bacterial pathogen *Pseudomonas syringae* pathovar *maculicola* ES4326 (*Psm* ES4326) (25.8%; 237/917 genes)^1, 2, 23–25^ (Fig. 2a, b). In total, 58.6% (537/917 genes) of raindrop- and MS-induced genes overlapped with those induced in response to different immune elicitors, suggesting that raindrops activate mechanosensitive immune responses.

**Fig. 2.**
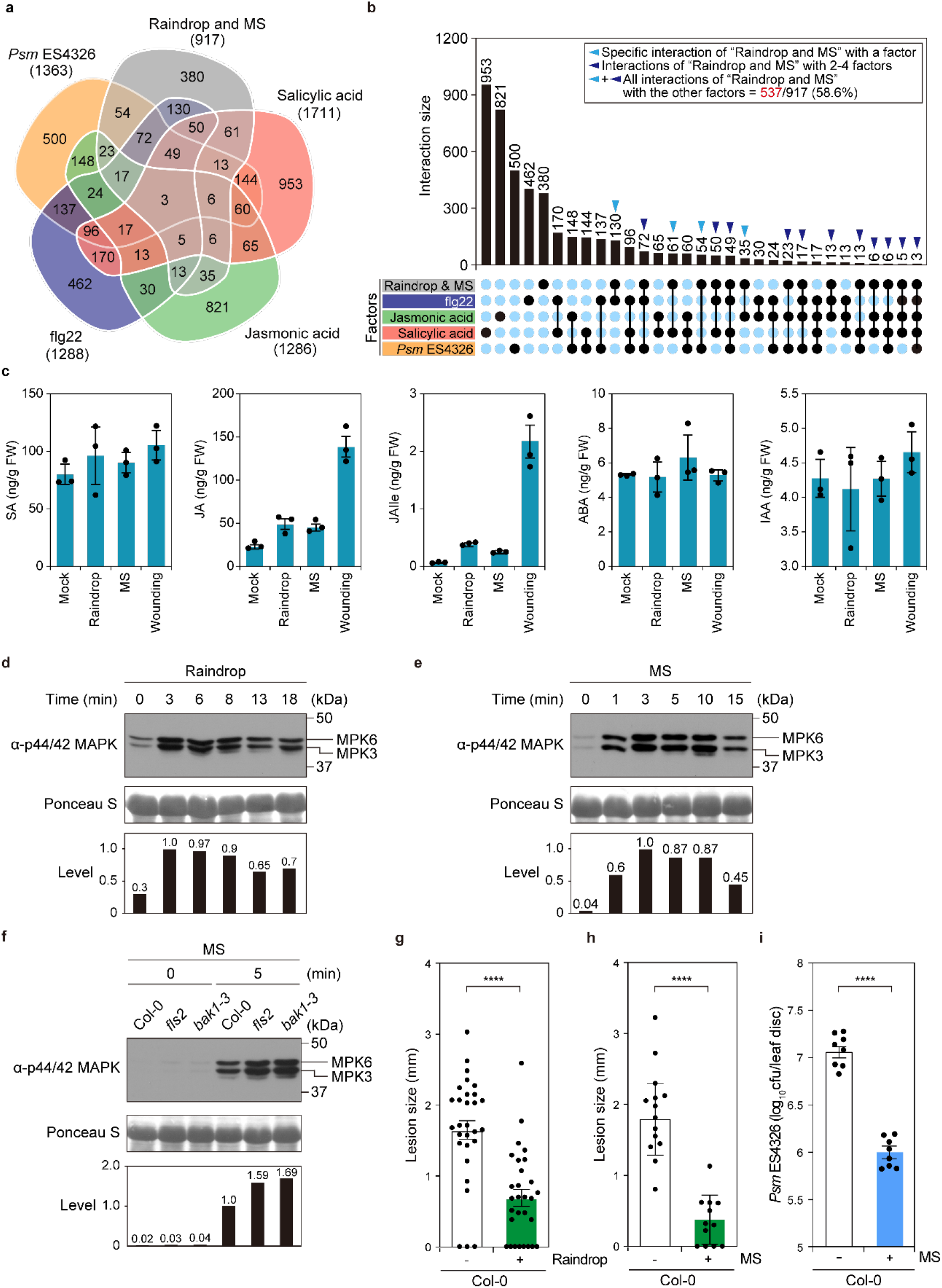
Raindrop-induced mechanosensation triggers defence responses in *Arabidopsis.* **a**, **b** Venn diagram (**a**) and the upset plot (**b**) between 917 raindrop- and MS-induced genes and transcriptome datasets obtained from salicylic acid (SA), jasmonic acid (JA), and flg22 (PAMP) treatment and *Psm* ES4326 infection (*P* < 0.05). Overlap with raindrop- and MS-induced genes: SA, 21%, 193/917 genes; JA, 11.8%, 108/917 genes; flg22, 37%, 339/917 genes; *Psm* ES4326, 25.8%, 237/917 genes; any of the four factors, 58.6%, 537/917 genes. **c** Fresh weight (ng/g) of plant hormones SA, JA, JA-isoleucine (JA-Ile), abscisic acid (ABA), and indole-3-acetic acid (IAA) 5 min after 10 falling droplets (raindrop), 1 brushing (MS), and cutting (wounding). **d**, **e** Raindrop (4 droplets)- (**d**) and MS (4 brushing)-induced (**e**) MAPK activation in Col-0. Total proteins were extracted from 4-week-old plants treated with raindrops and detected by immunoblot analysis with anti-p44/42 MAPK antibodies. Relative phosphorylation levels are shown below each blot. **f** MS-induced MAPK activation in Col-0, *fls2*, and *bak1-3*. Total proteins were extracted from 4-week-old plants after 5 min of MS treatment (1 brushing) and detected by immunoblot analysis with anti-p44/42 MAPK antibodies. Relative phosphorylation levels are shown below each blot. **g, h** Disease progression of *Alternaria brassicicola* in Col-0 leaves 3 days after inoculation with (+) or without (–) raindrop (10 droplets) pretreatment (**g**) or with (+) or without (–) MS (4 brushing) pretreatment (**h**). Error bars represent SE. Asterisks indicate significant difference (one-way ANOVA; \*\*\*\**P* < 0.0001). **i** Growth of *Psm* ES4326 in Col-0 leaves 2 days after inoculation with (+) or without (–) MS (4 brushing) pretreatment. An outline of the experiment is provided at left. Error bars represent SE. Asterisks indicates significant difference (one-way ANOVA; \*\*\*\**P* < 0.0001). Cfu, colony-forming units.

Since stress-responsive gene expression is either positively or negatively regulated by phytohormones, we determined the changes in the accumulation levels of six phytohormones [SA, JA, JA-isoleucine (JA-Ile), abscisic acid (ABA), gibberellic acid 4 (GA_4_), and indole-3-acetic acid (IAA)] in leaves treated with 10 falling droplets and in those brushed once. No significant changes in the levels of the phytohormones, except JA and JA-Ile, were observed 5 min after treatment (Fig. 2c), consistent with the previous report that water spray induces JA-mediated transcriptional changes^20^. The slight increase in JA and JA-Ile could explain the observation that only 11.8% of raindrop- and MS-induced genes are JA-responsive (Fig. 2a). Although 21% of raindrop- and MS-induced genes overlap with SA-responsive genes (Fig. 2a), SA levels were not significantly increased in response to raindrops and MS (Fig. 2c). Therefore, most mechanosensitive genes, whose expression is induced 5 min after treatment with raindrops, are presumably regulated independently of phytohormonal responses. A previous report demonstrated that GA accumulation is reduced by “bending” leaves twice per day for 2 weeks^19^. Here, significant changes in GA levels were not detected upon transient application of raindrops or MS (data not shown). These results indicated that plants differentially respond to MS depending on their intensity and duration.

Activation of MAPKs is one of the earliest cellular events and a hallmark of plant immune responses. In particular, PRRs promptly activate a phosphorylation cascade involving MPK3 and MPK6 in response to MAMPs, whereby the downstream immune components of PTI are phosphorylated to promote transcriptional reprogramming^1, 6, 7, 26^. Because 37% of raindrop- and MS-induced genes were also upregulated by flg22 treatment (Fig. 2a), we examined whether a MAPK cascade is activated in responses to raindrops and MS by immunoblot analysis with the anti-p44/p42 antibody, which detects phosphorylated MPK3/MPK6^27, 28^. Upon treatment of rosette leaves with 4 falling raindrops or MS (1 brushing), phosphorylation of MPK3/MPK6 was induced within 3 min and remained high for 10 min after each treatment (Fig. 2d, e), indicating that MPK3/MPK6 activation precedes the expression of mechanosensitive genes detected 10 min after MS application (Supplementary Fig. 1d). The kinetics of MS-activated MPK3/MPK6 were reminiscent of those observed upon activation of the PRR protein FLS2 and its coreceptor BRI1-ASSOCIATED RECEPTOR KINASE 1 (BAK1), which are responsible for recognition of the bacterial flg22 epitope^1, 26^. Wild-type, *fls2*, and *bak1* mutant plants displayed comparable levels of phosphorylated MPK3/MPK6 in response to MS (Fig. 2f), however, suggesting that FLS2 and BAK1 are not positively involved in raindrop-elicited mechanotransduction.

We then performed a comparative analysis of raindrop- and MS-induced genes against published transcriptome datasets describing the specific and conditional activation of MPK3/MPK6 in transgenic *Arabidopsis* plants carrying a constitutively active variant of tobacco (*Nicotiana tabacum*) *MAP KINASE Cab 2* (*NtMEK2*) under the control of the dexamethasone-inducible promoter^28^. Approximately 27.5% (252/917 genes) of both raindrop- and MS-induced genes were upregulated by MPK3/MPK6^28^ (Supplementary Fig. 2a, Supplementary Table 5), and these upregulated genes were highly enriched in categories associated with plant immunity (Supplementary Fig. 2b, Supplementary Table 6), suggesting that MAPKs play a critical role in mechanotransduction.

### Rain and MS confer resistance to both biotrophic and necrotrophic pathogens

We then investigated whether raindrops and MS confer resistance to pathogenic microbes. Raindrops containing the spores of the necrotrophic pathogen *Alternaria brassicicola* Ryo-1 were placed on fully expanded leaves after pretreatment with raindrops or MS for 3 h at an interval of 15 min. Both stimuli significantly suppressed lesion development compared to control plants without pretreatment (Fig. 2g, h). Pretreatment of leaves with MS for 3 h also efficiently protected plants from infection with the biotrophic pathogen *Psm* ES4326 (Fig. 2i). These results confirmed that mechanostimulation induces a PTI-like response to confer a broad spectrum of resistance to both biotrophic and necrotrophic pathogens, as MS activate immune MAPKs and upregulate a large subset of flg22-induced genes. In support of this argument, exposure to the fungal cell wall, chitin, also upregulated 42.1% (386/917 genes) of raindrop-induced genes (Supplementary Fig. 2c).

### Mechanosensitive genes are regulated by calmodulin-binding transcription activator 3

To dissect rain-induced mechanotransduction, we searched for a conserved *cis*-regulatory element in the promoter sequences of mechanosensitive genes. From an unbiased promoter analysis of the top 300 genes among 917 differentially expressed genes, we obtained the highest enrichment for the CGCG box (CGCGT or CGTGT), which is recognized by calmodulin (CaM)-binding transcription activators (CAMTAs) that are conserved from plants to mammals^29–33^ (Fig. 3a). The *Arabidopsis* transcription factor CAMTA3 (also named SIGNAL RESPONSIVE1 [SR1]) is a negative regulator of plant immunity because *camta3* null mutants exhibit constitutive expression of defence-related genes and enhanced resistance to virulent *P*. *syringae* infection^34, 35^. CAMTA transcription factors possess a CaM-binding (CaMB) domain and an IQ domain to which CaM binds in a calcium-dependent manner to negate their function (Supplementary Fig. 3a). Mutation of the IQ domain, such as in CAMTA3^A855V^, suppresses the constitutive expression of defence-related genes seen in the *camta2 camta3* double mutant when expressed in this background but is no longer regulated by calcium-mediated responses^36, 37^. In agreement with our promoter analysis, 28.7% of constitutively upregulated genes (309/1,075 genes) in the *camta1 camta2 camta3* triple mutant overlapped with raindrop- and MS-induced genes detected in wild-type plants^38^ (Supplementary Fig. 3b, Supplementary Table 7). Upon application of raindrops and MS, *WRKY33* and *CBP60g* transcript levels were significantly reduced in plants expressing the *CAMTA3^A855V^* variant compared to a *CAMTA3-GFP* transgenic line expressing a transgene that complemented the phenotype of the *camta2 camta3* mutant (Fig. 3b, c), suggesting that CAMTA3 is involved in mechanotransduction.

**Fig. 3.**
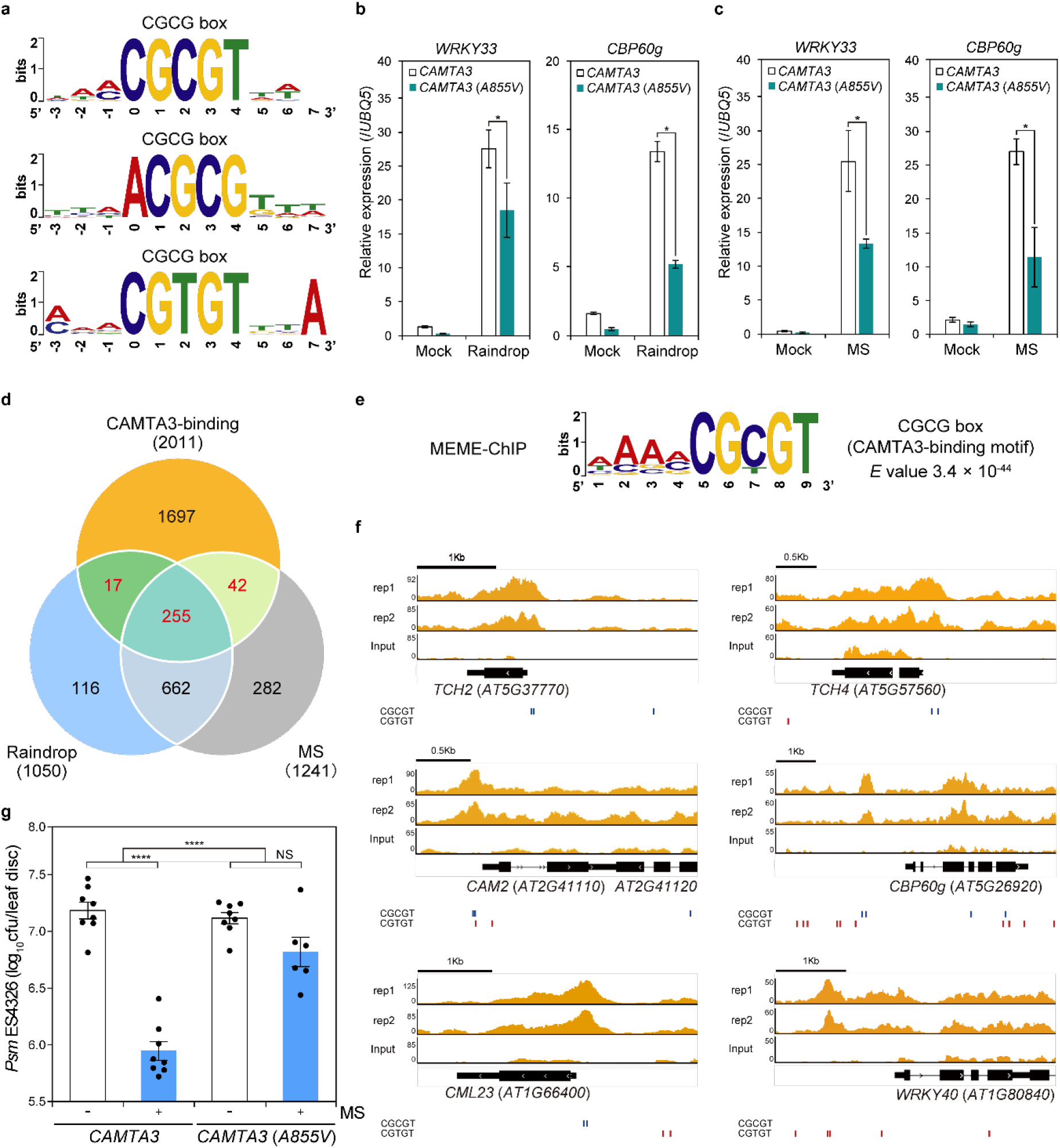
MS-induced genes are regulated by CAMTA3. **a** Promoter analysis of the top 300 (among 917 genes) raindrop- and MS-induced genes in terms of expression levels revealed that the CAMTA-binding CGCG box [CGC(/T)GT] were overrepresented among these genes. **b**, **c** Transcript levels of *WRKY33* and *CBP60g* in 4-week-old *camta2 camta3 CAMTA3pro:CAMTA3-GFP* (*CAMTA3*) and *camta2 camta3 CAMTA3pro:CAMTA3^A855V^-GFP* [*CAMTA3*(*A855V*)] plants 15 min after the plants were treated with 1 falling droplet (**b**) or brushed 4 times (**c**), determined by RT-qPCR and normalized to *UBQ5* transcript levels. Data are presented as mean ± SD. **d** Venn diagram depicting the overlap between genes with CAMTA3-binding sites in their promoters, as determined by ChIP-seq, and raindrop- and MS-induced genes as determined by RNA-seq. A total of 314 genes, shown in red, were identified as CAMTA3-target genes. **e** The CGCG box was identified as an overrepresented motif among the sequence peaks of 314 genes by MEME-ChIP. **f** Localization of CAMTA3 on the promoters of the MS-induced genes *TCH2*, *TCH4*, *CAM2*, *CBP60g*, *CML23*, and *WRKY40*, as representative of the 314 genes shown in (**d**). Blue and red lines indicate CGCGT and CGTGT, respectively. **g** Growth of *Psm* ES4326 in *camta2 camta3 CAMTA3pro:CAMTA3-GFP* (*CAMTA3*) and *camta2 camta3 CAMTA3pro:CAMTA3^A855V^-GFP* [*CAMTA3*(*A855V*)] plants 2 days after inoculation with (+) or without (–) MS (4 brushing) pretreatment. Error bars represent SE. Asterisks indicate significant difference (one- and two-way ANOVA; \*\*\*\**P* < 0.0001). Cfu, colony-forming units. NS, not significant.

To confirm whether CAMTA3 directly targets mechanosensitive genes, we investigated the genome-wide distribution of CAMTA3-binding sites by chromatin immunoprecipitation followed by deep sequencing (ChIP-seq) using *CAMTA3^A855V^-GFP* plants, as the mutant protein stably represses the transcription of CAMTA3-regulated genes. With the aid of model-based analysis of ChIP-seq (MACS2) software, we identified 2,641 and 2,728 CAMTA3-binding genes, respectively, in two replicates (*P* < 0.05); about 40% of these peaks are located in the promoter regions and another 30% in gene bodies (Supplementary Fig. 3c, Supplementary Table 8). The overlap between the two replicates highlighted 2,011 CAMTA3-targeted genes that included 272 raindrop- and 297 MS-induced genes such as *TCH2*, *TCH4*, and *CBP60g* (Fig. 3d), consistent with our hypothesis that CAMTA3 regulates the transcription of mechanosensitive genes.

To validate the results from the promoter analysis of mechanosensitive genes, we next investigated specific DNA sequences to which CAMTA3 selectively binds by analysing CAMTA3-binding peaks by Multiple EM for Motif Elicitation (MEME)-ChIP (Methods). We again identified the CGCG box (CGCGT or CGTGT) as the motif with the highest enrichment score (3.4 x 10^-44^) (Fig. 3e). The subsequent visualization of ChIP-seq profiles via the Integrative Genomics Viewer (IGV)^39^ demonstrated that CAMTA3 is primarily enriched at the CGCG boxes of mechanosensitive genes, including *TCH2*, *TCH4*, *CAM2*, *CBP60g*, *CALMODULIN LIKE 23* (*CML23*), and *WRKY40* (Fig. 3f). GO analysis on 314 CAMTA-targeted genes (Fig. 3d, shown in red) to define the biological functions of these genes showed a significant enrichment in categories related to immune and environmental responses (Supplementary Fig. 3d, e). We thus investigated whether CAMTA3 is required for the immune responses. *camta2 camta3 CAMTA3-GFP* transgenic plants effectively mounted an enhanced disease resistance against *P*. *syringae* in response to MS, while *camta2 camta3 CAMTA3^A855V^-GFP* plants were significantly compromised in resistance (Fig. 3g). These results demonstrate that CAMTA3 negatively regulates the plant immune responses by binding to the CGCG box in raindrop- and MS-induced gene promoters and represses the expression of these genes.

Since mechanostimulation rapidly activates MPK3/MPK6 (Fig. 2d, e), we investigated whether CAMTA3 mediates the activation of these MPKs. Using *camta2 camta3 CAMTA3-GFP* and *camta2 camta3 CAMTA3^A855V^-GFP*, we detected the phosphorylation of MPK3/MPK6 independently of CAMTA3 activity (Supplementary Fig. 3f). In addition, the calcium ionophore A23187 clearly induced the phosphorylation of MPK3 and MPK6 (Supplementary Fig. 3g). These results suggested that the mechanotransduction initiated by raindrops and MS may cause a Ca^2+^ influx that negates the repressive effect of CAMTA3 and concomitantly activates the MAPK cascade, as previously proposed^26^.

### MS initiates intercellular calcium waves concentrically away from trichomes

To visualize how mechanostimulation induces the expression of immune genes *in planta*, we generated *Arabidopsis* transgenic lines with the promoter sequences of *WRKY33* and *CBP60g* driving the expression of nucleus-targeted enhanced *YELLOW FLUORESCENT PROTEIN* (*YFP-NLS*) (*WRKY33pro:EYFP-NLS* and *CBP60gpro:EYFP-NLS*). *WRKY33* expression is regulated by both MPK3/MPK6 and CAMTA3 (Supplementary Table 5, Supplementary Table 8), while *CBP60g* is not mediated by MPK3/MPK6 (Supplementary Table 5). When half leaves were gently brushed (Supplementary Fig. 1c), we detected YFP fluorescence in the *WRKY33pro:EYFP-NLS* and *CBP60gpro:EYFP-NLS* transgenic lines as localized, clustered groups of cells only in the brushed half (Fig. 4a, Supplementary Fig. 4a). Closer inspection of the stimulated regions revealed that both genes were induced in cells surrounding trichomes, hair-like structures projecting outward from the epidermal surface (Fig. 4b, c, Supplementary Fig. 4b, c).

**Fig. 4.**
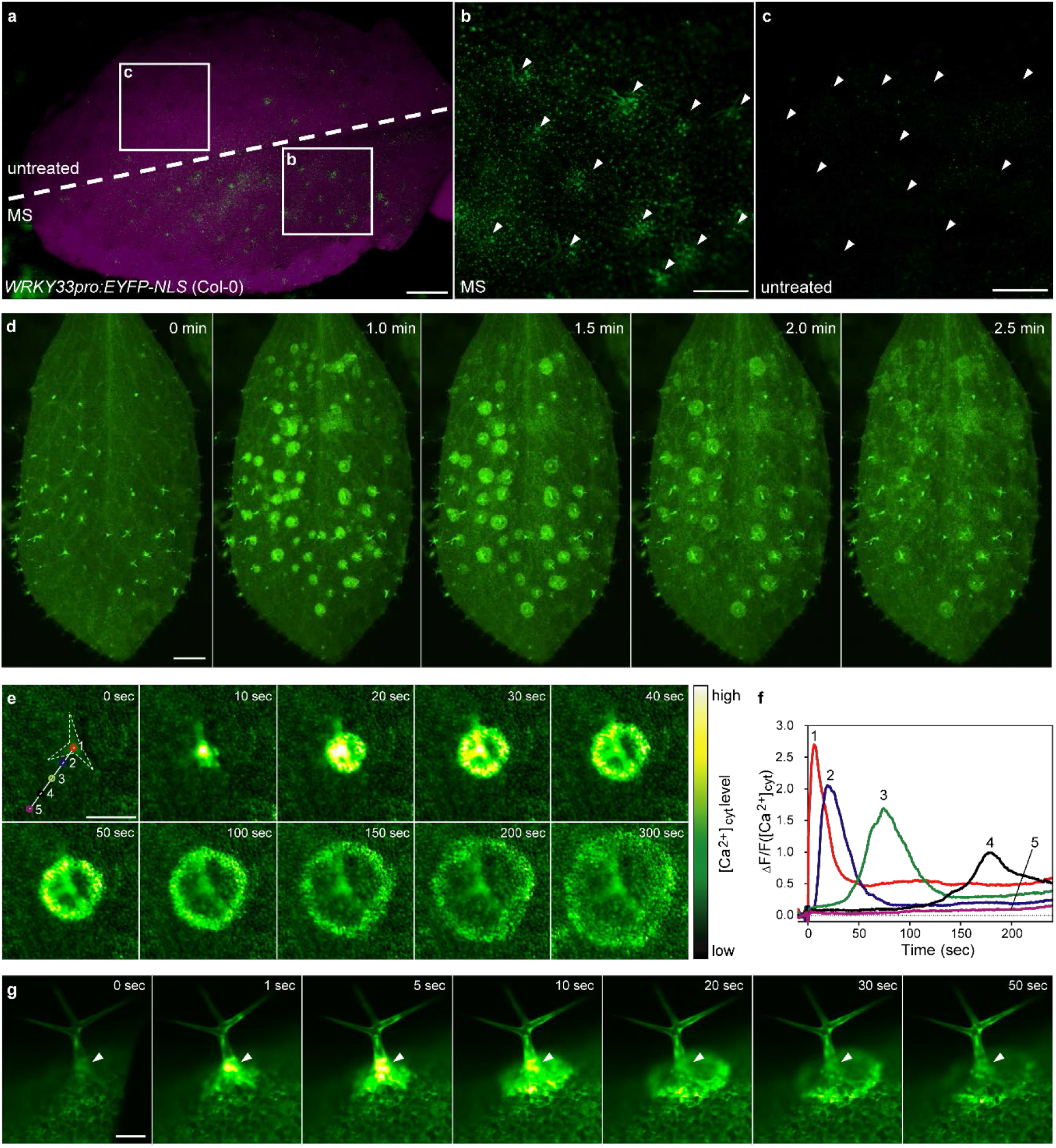
Trichomes initiate intercellular calcium waves. **a–c** YFP fluorescence in a whole leaf from *WRKY33pro:EYFP-NLS* (Col-0) with (MS, bottom half) or without brushing (untreated, top half) (**a**), along with zoomed-in views of brushed (**b**) and untreated (**c**) areas. Arrowheads indicate trichomes (**b, c**). Scale bars, 0.5 mm (**a**), 0.3 mm (**b**, **c**). **d** Ca^2+^ imaging using *35Spro:GCaMP3* (Col-0). The leaf surface of a 4-week-old plant was treated with MS by brushing. MS-induced intercellular calcium waves propagated concentrically from trichomes. Scale bar, 1.0 mm. See also Supplementary Video 1. **e** Ca^2+^ imaging using *35Spro:GCaMP3* (Col-0). A single trichome from a 2-week-old seedling was flicked with a silver chloride wire. MS-induced intercellular calcium waves propagated concentrically from the trichome (dashed outline). Scale bar, 0.2 mm. See also Supplementary Video 2. **f** [Ca^2+^]_cyt_ changes at sites indicated by numbers in (**e**). **g** Side view of a trichome whose neck was flicked with a silver chloride wire. MS-induced intercellular Ca^2+^ influx was transiently observed in the trichome base (arrowheads) followed by the formation of circular waves. Scale bar, 0.1 mm. See also Supplementary Video 3.

Trichomes function as chemical and physical barriers against insect feeding and are likely involved in drought tolerance and protection against ultraviolet irradiation^40, 41^. Mechanostimulation of a single trichome induces Ca^2+^ oscillations within the proximal skirt cells that surround the base of trichomes^42^, suggesting that the mechanical force could be focused on only skirt cells (Supplementary Fig. 5). However, since mechanostimulation by raindrops and MS confers resistance to pathogens in whole leaves, we hypothesized that trichomes activate a Ca^2+^ signal in a large area of leaves, as shown in Figure 4a.

To visualize changes in cytosolic Ca^2+^ concentrations ([Ca^2+^]_cyt_) induced by MS on the leaf surface, we used transgenic *Arabidopsis* expressing the GFP-based [Ca^2+^]_cyt_ indicator GCaMP3^43, 44^. Leaf brushing induced a marked increase of [Ca^2+^]_cyt_ in the surrounding leaf area of trichomes 1 min after stimulation (Fig. 4d, Supplementary Video 1). Flicking a single trichome with a silver chloride wire triggered an intercellular calcium wave that propagated concentrically away from the trichome and surrounding skirt cells at a speed of 1.0 μm/s (Fig. 4e, f, Supplementary Video 2). This pattern showed striking consistency with the area of induction observed with the *WRKY33pro:EYFP-NLS* and *CBP60gpro:EYFP-NLS* reporters (Fig. 4a, b; Supplementary Fig. 4a, b). The base of trichomes exhibited a rapid and transient increase in [Ca^2+^]_cyt_ before the concentric propagation of calcium waves was initiated (Fig. 4g, Supplementary Video 3).

### Trichomes are mechanosensory cells activating plant immunity

To investigate the possible involvement of trichomes in mechanosensation in *Arabidopsis* leaves and activation of the immune response, we observed calcium waves using the knockout mutant of *GLABROUS1* (*GL1*)^45^, which lacks trichomes. The *gl1* mutant exhibits effective basal resistance comparable to that of wild-type Col-0 plants^46^, and its local resistance to *Psm* ES4326 and *A. brassicicola* Ryo-1 is similar to the levels of Col-0 plants (Supplementary Fig. 6). The mechanostimulation-induced propagation of concentric calcium waves was compromised in the *gl1* mutant (Fig. 5a, b), confirming that trichomes are true MS sensors and initiate calcium waves (Supplementary Videos 4, 5). Furthermore, approximately 70.5% of mechanosensitive genes were expressed in a trichome-dependent manner (Fig. 5c, Supplementary Fig. 7a, Supplementary Table 9), and transcript levels of 18 representative mechanosensitive immune genes were markedly lower at all time points in the *gl1* mutant than they were in the wild type in RNA-seq analysis of leaves brushed 4 times (Fig. 5d), suggesting that trichomes serve as the main sensor of MS. Similarly, compared to wild-type plants, the transcription of raindrop-induced *WRKY33*, *WRKY40*, and *CBP60g*, as well as the activation of MPK3/MPK6 by MS, was also significantly reduced in the *gl1* mutant (Fig. 5e, f, Supplementary Fig. 7b). Moreover, MS-induced resistance against *Psm* ES4326 infection was abrogated in the *gl1* mutant (Fig. 5g). As with *Psm* ES4326, the application of MS to wild-type plants prior to inoculation with *A. brassicicola* significantly limited lesion development, whereas the *gl1* mutant did not show enhanced disease resistance in response to MS (Fig. 5h).

**Fig. 5.**
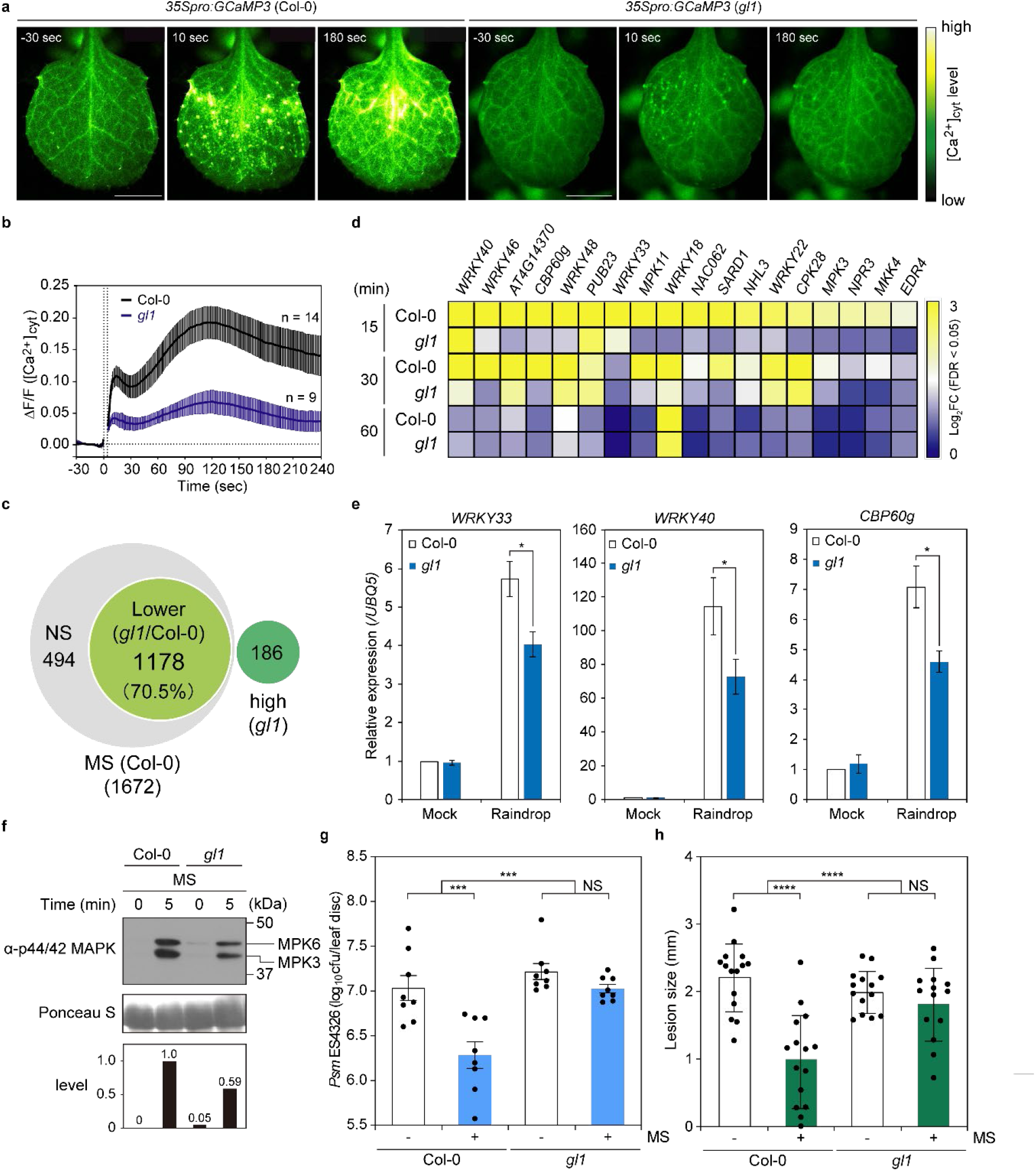
Trichomes are mechanosensory cells. **a** Ca^2+^ imaging using *35Spro:GCaMP3* (Col-0) and *35Spro:GCaMP3* (*gl1*). Leaf surfaces were exposed to MS by brushing. MS-induced calcium waves were compromised in the *gl1* mutant. See also Supplementary Videos 4 and 5. Scale bars, 0.5 mm. **b** [Ca^2+^]_cyt_ signature of (**a**). **c** Venn diagram of transcriptome datasets for MS-induced genes in Col-0 and *gl1* (*P* < 0.05). NS, not significant. Lower, fold change (FC) (*gl1*)/FC (Col-0) < 0.5. High, MS (*gl1*)/Mock (*gl1*), log_2_FC ≥ 1 in *gl1*. *P* < 0.05. **d** Heatmap of differentially expressed defence-related genes obtained from transcriptome datasets from Col-0 and *gl1* plants treated with MS (4 brushing). **e** Transcript levels of *WRKY33*, *WRKY40*, and *CBP60g* in 4-week-old Col-0 and *gl1* plants 15 min after treatment with 4 falling droplets, determined using RT-qPCR and normalized to *UBQ5*. Data are presented as mean ± SD. Asterisks indicate significant difference (one- and two-way ANOVA; \**P* < 0.05). **f** MS-induced MAPK activation in Col-0 and *gl1*. Total proteins were extracted from 4-week-old leaves 5 min after MS treatment and detected by immunoblot analysis with anti-p44/42 MAPK antibodies. Relative phosphorylation levels are shown below each blot. **g** Growth of *Psm* ES4326 in Col-0 and *gl1* leaves 2 days after inoculation with (+) or without (–) MS (4 brushing) pretreatment. Error bars represent SE. Asterisks indicate significant difference (one- and two-way ANOVA; \*\*\**P* < 0.001). Cfu, colony-forming units. NS, not significant. **h** Disease progression of *Alternaria brassicicola* in Col-0 and *gl1* leaves 3 days after inoculation with (+) or without (–) MS (4 brushing) pretreatment. Error bars represent SE. Asterisks indicate significant difference (one- and two-way ANOVA; \*\*\*\**P* < 0.0001). NS, not significant.

Our work highlights a novel layer of plant immunity that is triggered by an unexpected function of trichomes as mechanosensory cells. When trichomes are mechanically stimulated, intercellular calcium waves are concentrically propagated away from the trichomes, followed by the activation of CAMTA3-dependent immune responses (Fig. 6). Rapid phosphorylation of MAPKs also is a prerequisite for mechanosensitive gene expression, as MPK3/MPK6 mediate the phosphorylation of mechanosensitive WRKY33 for its activation^47, 48^. This notion is supported by the finding that the expression of 252 genes among 917 raindrop- and MS-induced genes is regulated by MPK3/MPK6 (Supplementary Fig. 2a), and their promoter sequences possess the W-box (TTGACC) for WRKYs as the most enriched *cis*-regulatory elements (Supplementary Fig. 8). The molecular mechanism by which calcium mediates the activation of MPK3/MPK6 has yet to be elucidated.

**Fig. 6.**
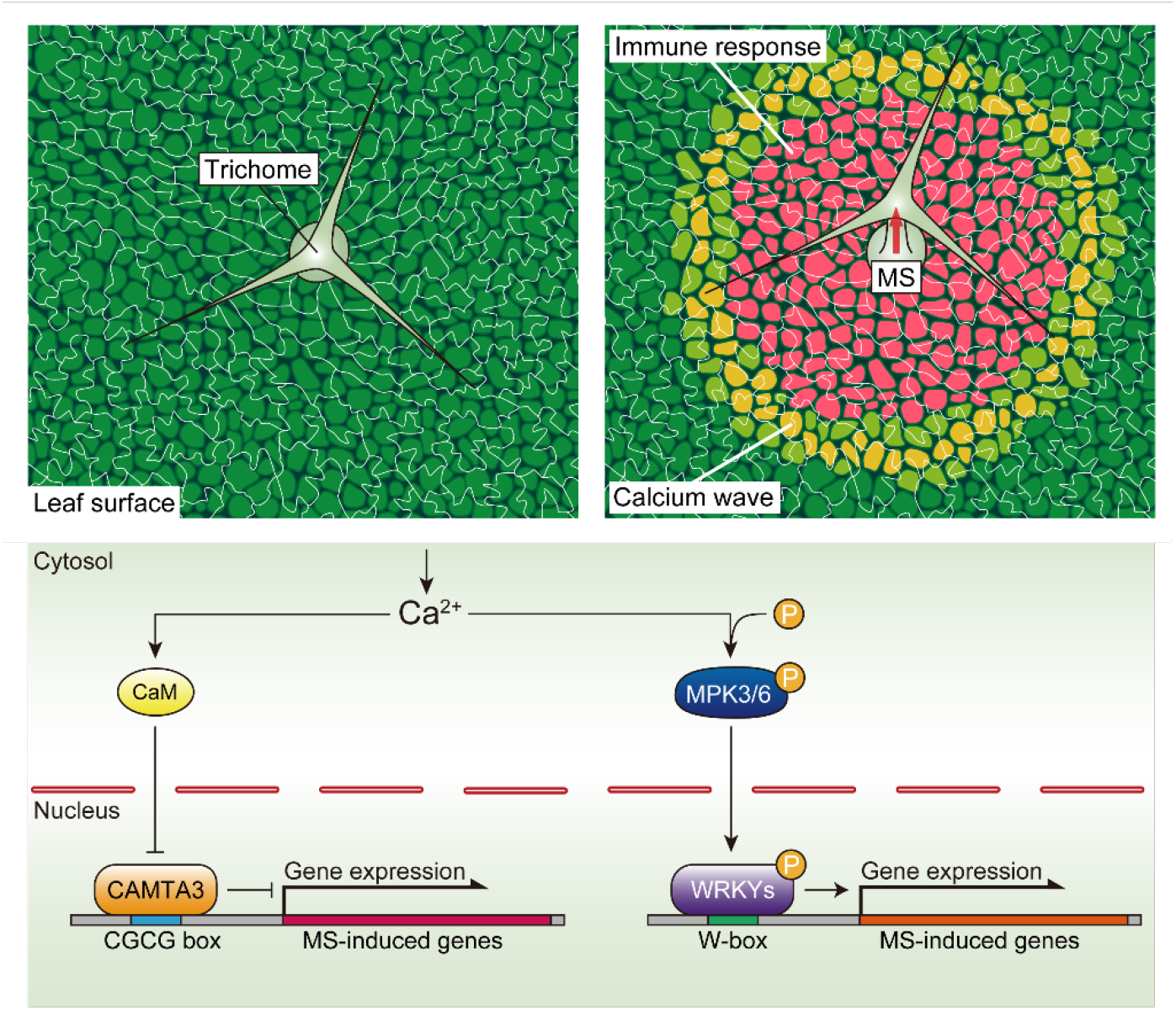
Trichomes activate broad-spectrum disease resistance. Model showing how trichomes directly sense the mechanical impact of raindrops as an emergency signal in anticipation of possible infections. Mechanosensory trichome cells initiate intercellular calcium waves in response to MS. [Ca^2+^]_cyt_ initiates the de-repression of Ca^2+^/CaM-dependent CAMTA3 and activates the phosphorylation of MPK3 and MPK6, thereby inducing WRKY-dependent transcription.

Mechanostimulation by repeatedly bending leaves confers resistance to the necrotrophic pathogen *Botrytis cinerea* via JA accumulation^49^. In addition, a subset of JA-responsive genes upregulated by water spray is mediated by MYC2/MYC3/MYC4 transcription factors^20^. These observations strongly indicate that mechanosensation causes profound JA-dependent changes in gene transcription, promoting plant immune responses to necrotrophic pathogens. The JA- and MYC-dependent pathway does not play a major role in the expression of mechanosensitive *TCH* genes, however, indicating that mechanotransduction is regulated by other signaling pathways. Our work demonstrated that raindrops and MS only partially activate the JA signal but rather strongly induce a PTI-like response via the Ca^2+^- and CAMTA3-dependent pathway, which is highly effective against both necrotrophs and biotrophs (Fig. 2g-i). Because rain disseminates diverse pathogens with different parasitic strategies, including fungi, bacteria, and virus^50, 51^, it is highly reasonable that plants perceive raindrops as a risk factor and activate broad-spectrum resistance.

Plants possess mechanosensory cells with a variety of functions, such as flower antennas of *Catasetum* species for pollination, tentacles of *Drosera rotundifolia* for insect trapping, root hairs of *Arabidopsis* for water tracking, and red cells of *Mimosa pudica* for evading herbivores^52^. The carnivorous Venus flytrap (*Dionaea muscipula*) captures insects by sensing mechanostimulation via sensory hairs on leaf lobes^53^. To monitor diverse MS applied to plants, several sensing mechanisms have been proposed that include the detection of cell wall components, distortion of the plasma membrane, and the displacement of the plasma membrane against the cell wall^54^. In all these systems, a transient increase in [Ca^2+^]_cyt_ is thought to play a pivotal role in short- and long-term responses. Indeed, two successive stimulations of sensory hairs of the flytrap are required to meet the threshold of [Ca^2+^]_cyt_ for rapid closure of the leaf blade^53, 55^. As the trichome on the leaf surface is widely conserved among many land plants, there may be a common and novel intercellular network of cell-cell communication that initiates calcium waves for activating immune responses.

## Contact for reagents and subject details

Further information and requests for reagents may be directed to and will be fulfilled by the corresponding authors Yasuomi Tada and Mika Nomoto.

## Experimental model and subject details

### Plants

*Arabidopsis thaliana* accession Columbia-0 (Col-0) was the background for all plants used in this study. *WRKY33pro:EYFP-NLS* (Col-0) and *CBP60gpro:EYFP-NLS* (Col-0) were generated as previously described^56^. *35Spro:GCaMP3* (Col-0), *gl1* [Col(*gl1*)], *camta2 camta3 CAMTA3pro:CAMTA3-GFP*, and *camta2 camta3 CAMTA3pro:CAMTA3^A855V^-GFP* were previously reported^36, 38^. The *Arabidopsis* mutants *fls2* (SALK_093905) and *bak1-3* (SALK_034523) were obtained from the Arabidopsis Biological Resource Center (ABRC). *35Spro:GCaMP3* was introduced into the *gl1* mutant background by crossing. The selection of homozygous lines was performed by genotyping using primers listed in Supplementary Table 10. Plants were grown on soil (peat moss; Super Mix A and vermiculite mixed 1:1) at 22°C under diurnal conditions (16-h-light/8-h-dark cycles) with 50-70% relative humidity. *WRKY33pro:EYFP-NLS* (Col-0) and *CBP60gpro:EYFP-NLS* (Col-0) were sown on soil and grown in a growth room at 23°C in constant light as previously described^56^. *35Spro*:*GCaMP3* (Col-0) and *35Spro*:*GCaMP3* (*gl1*) were grown on Murashige and Skoog (MS) plates [1% (w/v) sucrose, 0.01% (w/v) myoinositol, 0.05% (w/v) MES, and 0.5% (w/v) gellan gum pH 5.8] as previously described^44, 57^.

### Artificial raindrop treatment

Reverse osmosis (RO) water was kept in a 500 mL beaker until the water temperature reached room temperature (22°C). A transfusion set (NIPRO Infusion Set TI-U250P, Nipro, Osaka, Japan) was installed on a steel stand with the beaker at a height of 1.2 m (H-type Stand I3, As One, Osaka, Japan) and was adjusted to release 13 µL water droplets (Supplementary Fig. 1a). In this setting, the applied mechanical energy to the leaf surface is equivalent to one in which 5.8 µL of raindrops reach a terminal velocity of 6.96 m/s^58^. This size raindrop is frequently observed in nature; thus, the impact of simulated rain is comparable with that of true rain^58^. The adaxial side of leaves from 4-week-old plants was treated with 10 droplets for RNA-seq, and 1, 4 or 10 droplets for quantitative RT-PCR (RT-qPCR). The adaxial side of leaves from 4-week-old plants was treated with one falling or static droplet (Supplementary Fig. 1b). Sample leaves were collected 15 min after treatment and stored at −80 °C until use.

### Brush treatment

The adaxial side of leaves from 4-week-old plants was brushed once for RNA-seq and 4 for RT-qPCR along the main veins at an angle of 30-40° (KOWA nero nylon drawing pen flat 12, Kowa, Aichi, Japan) (Supplementary Fig. 1c). Sample leaves were collected 15-, 30- and 60 min after treatment for RNA-seq and 15 min for RT-qPCR, and stored at -80°C until use.

### RNA-seq library construction

Total RNA was extracted from 80-100 mg frozen samples using Sepasol-RNA I Super G (Nacalai Tesque, Kyoto, Japan) and the TURBO DNase free kit (Thermo Fisher Scientific, IL, USA) according to the manufacturers’ protocols. Total RNA was further purified with the RNeasy RNA Isolation Kit (QIAGEN, Hilden, Germany) and assessed for quality and quantity with a Nanodrop spectrophotometer (Thermo Fisher Scientific). We used 1 µg total RNA for mRNA purification with NEBNext Oligo d(T)_25_ (NEBNext poly(A) mRNA Magnetic Isolation Module; New England Biolabs, MA, USA), followed by first-strand cDNA synthesis with the NEBNext Ultra DNA Library Prep Kit for Illumina (New England Biolabs) and NEBNext Multiplex Oligo for Illumina (New England Biolabs) according to the manufacturer’s protocols. For the analysis of raindrop- and MS-induced gene expression, the amount of cDNA was determined on an Agilent 4150 TapeStation System (Agilent, CA, USA). cDNA libraries were sequenced as single-end reads for 81 nucleotides on an Illumina Nextseq 550 (Illumina, CA, USA). The reads were mapped to the *Arabidopsis thaliana* reference genome (TAIR10, http://www.arabidopsis.org/) online (BaseSpace, Illumina, https://basespace.illumina.com/). Pairwise comparisons between samples were performed with the EdgeR^59^ package on the web (Degust, https://degust.erc.monash.edu/). For the comparative analysis of differentially expressed genes between leaves in the *gl1* mutant and Col-0, the amount of cDNA was determined by the QuantiFluor dsDNA System (Promega, WI, USA). cDNA libraries were sequenced as single-end reads for 36 nucleotides on an Illumina Nextseq 500 (Illumina). The reads were mapped to the *Arabidopsis thaliana* reference genome (TAIR10) via Bowtie^60^ with the options “--all --best --strata”. Pairwise comparisons between samples were performed with the EdgeR package in the R program^59^. Enrichment of GO categories for biological processes was determined using BiNGO (http://www.psb.ugent.be/cbd/papers/BiNGO/Home.html) (*P* < 0.05)^61^.

### Re-analysis of immune-related transcriptome datasets

We used the following public transcriptome datasets for the comparative analysis with the RNA-seq data obtained in this study: 10-day-old *Arabidopsis* seedlings treated with 1 µM flg22 (Array Express; E-NASC-76)^62^, 8-day-old *Arabidopsis* seedlings treated with 40 µM chitin (Gene Expression Omnibus GSE74955), leaves from 4-week-old *Arabidopsis* plants inoculated with *Pseudomonas syringae* pv. *maculicola* (*Psm*) ES4326 (24 h post inoculation) (GSE18978), 2-week-old *Arabidopsis* seedlings treated with 0.5 mM SA or 50 µM JA (DNA Data Bank of Japan DRA003119), 12-day-old *Arabidopsis GVG-NtMEK2^DD^* seedlings treated with 2 μM DEX for 0 and 6 h (NCBI Sequence Read Archive SRP111959), and 4-week-old *Arabidopsis camta1 camta2 camta3* triple mutant (GSE43818). The overlaps between differentially expressed genes in each transcriptome dataset were evaluated as Venn diagrams (http://bioinformatics.psb.ugent.be/webtools/Venn/).

### RT-qPCR

Total RNA was extracted from 30-40 mg leaf tissues with Sepasol-RNA I Super G and the TURBO DNase free kit (Thermo Fisher Scientific) according to the manufacturer’s protocols, followed by reverse transcription with the PrimeScript RT reagent kit (Takara Bio, Shiga, Japan) using oligo dT primers. RT-qPCR was performed on the first-strand cDNAs diluted 20-fold in water using KAPA SYBR FAST qPCR Master Mix (2x) kit (Roche, Basel, Switzerland) and gene-specific primers in a LightCycler 96 (Roche). Primer sequences are listed in Supplementary Table 10.

### Quantification of plant hormones

The adaxial side of leaves from 4-week-old plants was treated with 10 raindrops (raindrop), brushed once (MS), and cut (wounding). Sample leaves (0.07-0.1 g) were collected 5 min after treatment and stored at -80°C until use. SA, JA, JA-Ile, ABA, IAA, and GA_4_ were extracted and purified by solid-phase extraction. The contents of these hormones were quantified using liquid chromatography-electrospray tandem mass spectrometry (LC-ESI-MS/MS) (triple quadrupole mass spectrometer with 1260 high-performance LC, G6410B; Agilent Technologies Inc., CA, USA), as previously reported^63^.

### ChIP assay

Approximately 0.7 g of 2-week-old *camta2 camta3 CAMTA3pro:CAMTA3^A855V^-GFP* seedlings was fixed in 25 mL 1% formaldehyde under vacuum for three cycles of 2 min ON/2 min OFF using an aspirator (SIBATA, Tokyo, Japan). Subsequently, 1.5 mL of 2 M glycine was added to quench the cross-linking reaction under vacuum for 2 min. The samples were then washed with 50 mL double-distilled water and stored at -80°C until use. Frozen samples were ground to a fine powder with a mortar and pestle in liquid nitrogen and dissolved in 2.5 mL nuclei extraction buffer (10 mM Tris-HCl pH 8.0, 0.25 M sucrose, 10 mM MgCl_2_, 40 mM β-mercaptoethanol, protease inhibitor cocktail)^64^. Samples were filtered through two layers of Miracloth (Calbiochem, CA, USA) and centrifuged at 17,700 g at 4°C for 5 min. The pellets were resuspended in 75 μL nuclei lysis buffer [50 mM Tris-HCl pH 8.0, 10 mM EDTA, 1% (w/v) SDS]. After incubation first at room temperature for 20 min and then on ice for 10 min, the samples were mixed with 225 μL ChIP dilution buffer without Triton [16.7 mM Tris-HCl pH 8.0, 167 mM NaCl, 1.2 mM EDTA, 0.01% (w/v) SDS]. Chromatin samples were sonicated for 35 cycles of 30 sec ON/30 sec OFF using a Bioruptor UCW-201 (Cosmo Bio, Tokyo, Japan) to produce DNA fragments, followed by the addition of 375 μL ChIP dilution buffer without Triton, 200 μL ChIP dilution buffer with Triton [16.7 mM Tris-HCl pH 8.0, 167 mM NaCl, 1.2 mM EDTA, 0.01% (w/v) SDS, 1.1% (w/v) Triton X-100], and 35 μL 20% (w/v) Triton X-100. After centrifugation at 17,700 g at 4°C for 5 min, 900 μL solubilized sample was split into two 2.0 mL PROKEEP low-protein-binding tubes (Watson Bio Lab USA, CA, USA) and incubated with 0.75 μL anti-GFP antibody (for immunoprecipitation [IP]) (ab290; Abcam, Cambridge, UK) or Rabbit IgG-Isotype Control (Input) (ab37415; Abcam) for 4.5 h with gentle rocking, and an 18 μL aliquot was used as the input control. Then, samples from *camta2 camta3 CAMTA3pro:CAMTA3^A855V^-GFP* were mixed with 50 µL of a slurry of Protein A agarose beads (Upstate, Darmstadt, Germany) and incubated at 4°C for 1 h with gentle rocking. Beads were washed twice with 1 mL low-salt wash buffer [20 mM Tris-HCl pH 8.0, 150 mM NaCl, 2 mM EDTA, 0.1% (w/v) SDS, 1% (w/v) Triton X-100], twice with 1 mL high-salt wash buffer [20 mM Tris-HCl pH 8.0, 500 mM NaCl, 2 mM EDTA, 0.1% (w/v) SDS, 1% (w/v) Triton X-100], twice with 1 mL LiCl wash buffer [10 mM Tris-HCl pH 8.0, 0.25 M LiCl, 1 mM EDTA, 1% (w/v) sodium deoxycholate, 1% (w/v) Nonidet P-40], and twice with 1 mL TE buffer [10 mM Tris-HCl pH 8.0, 1 mM EDTA]. After washing, beads were resuspended in 100 μL elution buffer [1% (w/v) SDS, 0.1 M NaHCO_3_] and incubated at 65°C for 30 min. For the input controls, 41.1 μL TE buffer, 8.7 μL 10% (w/v) SDS, and 21 μL elution buffer were added to 18 μL of each solubilized sample. Both supernatant and input samples were mixed with 4 μL of 5 M NaCl and incubated at 65°C overnight to reverse the cross-linking, followed by digestion with 1 μL Proteinase K (20 mg/ml) (Invitrogen, CA, USA) at 37°C for 1 h. ChIP samples were mixed with 500 μL Buffer NTB and purified using the PCR clean-up gel extraction kit following the manufacturer’s instructions (MACHEREY-NAGEL, Düren, Germany).

### ChIP-seq library construction

ChIP-seq libraries for the input and two biological replicates were constructed from 2 ng purified DNA samples with the NEB Ultra II DNA Library Prep Kit for Illumina (New England Biolabs) according to the manufacturer’s instructions. The amount of DNA was determined on an Agilent 4150 TapeStation System (Agilent). All ChIP-seq libraries were sequenced as 81-nucleotide single-end reads using an Illumina NextSeq 550 system.

### Analysis of ChIP-seq

Reads were mapped to the *Arabidopsis thaliana* reference genome (TAIR10, http://www.arabidopsis.org/) using Bowtie2 with default parameters^60^. The Sequence Alignment/Map (SAM) file generated by Bowtie2 was converted to a Binary Alignment/Map (BAM) format file by SAMtools^65^. To visualize mapped reads, Tiled Data Files (TDF) file were generated from each BAM file using the igvtools package in the Integrative Genome Browser (IGV)^39^. ChIP-seq peaks were called by comparing the IP with the Input using Model-based Analysis of ChIP-Seq (MACS2) with the “-p 0.05 -g 1.19e8” option (*P* < 0.05)^66^. The peaks were annotated using the nearest gene using the Bioconductor and the ChIPpeakAnno packages in the R program, from which we identified 2,011 genes detected in both biological replicates. Enrichment of GO categories of the set of 314 genes overlapping between raindrop- and MS-induced genes for biological processes was determined using BiNGO (http://www.psb.ugent.be/cbd/papers/BiNGO/Home.html)61. Sequences of the peaks were extracted from the *Arabidopsis thaliana* genome as FASTA files with Bedtools^67^. To identify the candidates of CAMTA3-binding motifs, the FASTA files were subjected to MEME (Multiple EM for Motif Elicitation)-ChIP with the default parameter (-meme-minw 6-meme-maxw 10)^68^, and a density plot of the distribution of the motifs was generated.

### Immunoblot analysis for detection of MPK3 and MPK6 phosphorylation

The adaxial side of leaves from 4-week-old plants was brushed four times or treated with four raindrops, and samples (0.1-0.15 g) were snap-frozen in liquid nitrogen. Total proteins were extracted in protein extraction buffer [50 mM Tris-HCl pH 7.5, 150 mM NaCl, 2 mM DTT, 2.5 mM NaF, 1.5 mM Na_3_VO_4_, 0.5% (w/v) Nonidet P-40, 50 mM β-glycerophosphate, and proteinase inhibitor cocktail] and centrifuged once at 6,000 g, 4°C, for 20 min and twice at 17,000 g, 4°C for 10 min. The supernatant was mixed with SDS sample buffer [50 mM Tris-HCl pH 6.8, 2% (w/v) SDS, 5% (w/v) glycerol, 0.02% (w/v) bromophenol blue, and 200 mM DTT] and heated at 70°C for 20 min. The protein samples were subjected to SDS-PAGE electrophoresis and transferred onto a nitrocellulose membrane (GE Healthcare, IL, USA). The membrane was incubated with an anti-phospho-p44/42 MAPK polyclonal antibody (Cell Signalling Technology, MA, USA) (1:1,000 dilution) and goat anti-rabbit IgG(H+L)-HRP secondary antibody (BIO-RAD, CA, USA) (1:2,000 dilution). The bands for MPK3/6 were visualized using chemiluminescence solution mixed 5:1 with ImmunoStar Zeta (FUJIFILM Wako Chemicals, Osaka, Japan) and SuperSignal West Dura Extended Duration Substrate (Thermo Fisher Scientific). The Rubisco bands were stained with Ponceau S (Merck Sharp & Dohme Corp., NJ, USA) as a loading control. The phosphorylation levels of MPK3 and MPK6 were quantified with the blot analysis plug-in in ImageJ (https://imagej.nih.gov/ij/).

### Treatment with the calcium ionophore A23187

Twelve-day-old Col-0 seedling was treated with 50 µM calcium ionophore A23187 for 15-, 30- and 60 min. Samples were processed for the phosphorylation of MPK3 and MPK6 as described in the “Immunoblot analysis for detection of MPK3 and MPK6 phosphorylation” section. The leaf tissue was stored at -80°C until use.

### Promoter-reporter imaging

The 3.0-kbp promoters for *WRKY33* and *CBP60g*, both of which covered the previously analyzed respective regulatory sequences, were amplified from Col-0 genomic DNA by PCR and cloned into the pENTR/D-TOPO vector (Invitrogen). The promoter regions were recombined using Gateway technology into the binary vector pBGYN. The resulting pBGYN-pWRKY33-YFP-NLS and pBGYN-pCBP60g-YFP-NLS vectors were introduced into *Agrobacterium tumefaciens* GV3101 (pMP90) and then into *Arabidopsis* Col-0 plants using the floral dip method. A representative homozygous line was selected for each construct for further detailed analyses.

Promoter-reporter imaging was performed using an MA205FA automated stereomicroscope (Leica Microsystems, Wetzlar, Germany) and DFC365FX CCD camera (Leica Microsystems) in 12-bit mode. Chlorophyll autofluorescence and YFP fluorescence were detected through Texas Red (TXR) (excitation 560/40 nm, extinction 610 nm) and YFP (excitation 510/20 nm, extinction 560/40 nm) filters (Leica Microsystems). To image fluorescence emanating from the *WRKY33pro:EYFP-NLS* (Col-0) and *CBP60gpro:EYFP-NLS* (Col-0) plants^56^, the leaves of 3-week-old *Arabidopsis* plants were brushed 10 times at an interval of 15 min for 2 h or left untreated.

### Promoter analysis

The statistical analysis for overrepresented transcriptional regulatory elements across transcriptome datasets described above was calculated using a prediction program as previously reported^32^. The *P* values were calculated using Statistical Motif Analysis in Promoter or Upstream Sequences (https://www.arabidopsis.org/tools/bulk/motiffinder/index.jsp). Figures of promoter motif sequences are generated with WebLogo (https://weblogo.berkeley.edu/logo.cgi).

### Real-time [Ca^2+^]_cyt_ imaging

We used 4-week-old and 3-week-old plants expressing the GFP-based cytosolic Ca^2+^ concentration ([Ca^2+^]_cyt_) indicator GCaMP3^43, 44^. To image the fluorescence from the GCaMP3 reporter (in Col-0 and *gl1*) in whole leaves, the adaxial sides of leaves from 4-week-old plants were brushed. To monitor the calcium waves propagating from trichomes, a single trichome from a 2-week-old seedling was flicked with a silver chloride wire. Samples were imaged with a motorized fluorescence stereomicroscope (SMZ-25; Nikon, Tokyo, Japan) equipped with a 1× objective lens (NA = 0.156, P2-SHR PLAN APO; Nikon) and an sCMOS camera (ORCA-Flash 4.0 V2; Hamamatsu Photonics, Shizuoka, Japan) as described^44^.

### Propidium iodide staining

A stock solution of 10 mM propidium iodide (PI) was prepared with phosphate-buffered saline (PBS). Rosette leaves of 4-week-old Col-0 plants were cut into 5 mm squares, floated in a glass petri dish with 20 µM PI solution, and incubated for 1 h at room temperature. Stained tissues were observed under the all-in-one fluorescence microscope (BZ-X800; KEYENCE CORPORATION, Osaka, Japan) equipped with a 20x objective lens (CFI S Plan Fluor LWD ADM 20XC, Nikon) and TRITC dichroic mirror (excitation 545/25 nm, extinction 605/70 nm) (KEYENCE).

### Bacterial infection

MS were applied to the adaxial leaf surface of 4-week-old plants by brushing 4 times at an interval of 15 min for 3 h. Sample leaves were then inoculated by infiltration, using a plastic syringe (Terumo Tuberculin Syringe 1 mL; TERUMO), with *Psm* ES4326 (OD_600_ = 0.001) resuspended in 10 mM MgCl_2_. Bacterial growth was measured 2 days after inoculation as described previously^69^.

### Fungal infection

*Alternaria brassicicola* strain Ryo-1 was cultured on 3.9% (w/v) potato dextrose agar plates (PDA; Becton, Dickinson and Company, NJ, USA) for 4-20 days at 28°C in the dark. After incubation of the agar plates for 3-7 days under ultraviolet C light, a conidial suspension of *A. brassicicola* was obtained by mixing with RO water^70^. The adaxial side of leaves from 4-week-old plants was treated with 10 droplets or MS by brushing at an interval of 15 min for 3 h, followed by spotting with 5 µL conidia suspension (2 x 10^5^ per mL) of *A. brassicicola* on the adaxial side of leaves. Inoculated plants were placed at 22°C under diurnal conditions (16-h-light/8-h-dark cycles) with 100% relative humidity. The lesion size of fungal infection was measured with ImageJ 3 days after inoculation.

### Statistics and reproducibility

GraphPad Prism 9 (GraphPad software, CA, USA) was used for all statistical analyses. Two-sided one-way analysis of variance (one-way ANOVA) or two-way analysis of variance (two-way ANOVA) was used for multiple comparisons. Unless stated otherwise, sample size n represents technical replicates. In RT-qPCR, n ≥ 3; in bacterial growth assays, n = 8; in real-time [Ca^2+^]_cyt_ imaging assays of *35Spro:GCaMP3* (Col-0) and *35Spro:GCaMP3* (*gl1*), n = 14 and 9, respectively; and in fungal disease propagation assays, n = 29 (Fig. 2g) and n = 15 (the other figures). All experiments were performed at least three times with similar results (biological replicates). In all graphs, asterisks indicate statistical significance tested by Student’s *t* test (two groups) or one/two-way ANOVA (multiple groups).

### Reporting summary

Further information on research design is available in the Nature Research Reporting Summary linked to this article.

### Date availability

The authors declare that all data supporting the findings of this study are available within this article and its Supplementary Information files. RNA-seq and ChIP-seq data have been deposited in the DDBJ Sequence Read Archive (DRA) at the DNA Data Bank (DDBJ; http://www.ddbj.nig.ac.jp/) through the accession numbers DRA011970, DRA009248 and DRA011123.

## Acknowledgements

We thank M. F. Thomashow for the seeds of CAMTA3 variants. This work was supported by JSPS KAKENHI Grant Numbers JP23120520, JP25120718, and JP18K19334 (to Y.T.), JP15H05955 (to Y.T. and T.K.), JP19H05363, and JP21H00366 (to M.N.).

## Author contributions

M.M., M.N., S.H.S., and Y.T. designed the research. M.M. and T.I. established the artificial rain device. M.M. optimized the protocols for artificial raindrop and brush treatment. M.M. and M.N. constructed the Illumina sequencing libraries for RNA-seq and ChIP-seq. T.S. performed RNA-seq and analysis. M.N. performed the ChIP-seq and analysis of CAMTA3. T.M. and I.M. performed the quantification of phytohormones. M.M., Y.H., and T.K. performed the detection of MPK3 and MPK6 phosphorylation. Y.A. and M.T. generated *35Spro*:*GCaMP3* (*gl1*) plants, and M.N., M.T., and Y.A. visualized real-time [Ca^2+^]_cyt_. M.I. and S.B. generated the transgenic lines *WRKY33pro:EYFP-NLS* (Col-0) and *CBP60gpro:EYFP-NLS* (Col-0). M.M., M.I., and S.B. performed promoter-reporter imaging. M.N. and M.M. performed the rest of the experiments. M.M., M.N., S.H.S., and Y.T. wrote the manuscript with input from all authors.

## References

1. Couto, D. & Zipfel, C. Regulation of pattern recognition receptor signalling in plants. Nat. Rev. Immunol. 16, 537–552 (2016).

2. Zipfel, C. et al. Bacterial disease resistance in *Arabidopsis* through flagellin perception. Nature 428, 764–767 (2004).

3. Chisholm, S. T., Coaker, G., Day, B. & Staskawicz, B. J. Host-microbe interactions: shaping the evolution of the plant immune response. Cell 124, 803–814 (2006).

4. Cui, H., Tsuda, K. & Parker, J. E. Effector-triggered immunity: from pathogen perception to robust defense. Annu. Rev. Plant Biol. 66, 487–511 (2015).

5. Jones, J. D. & Dangl, J. L. The plant immune system. Nature 444, 323–329 (2006).

6. Boudsocq, M. et al. Differential innate immune signalling via Ca(2+) sensor protein kinases. Nature 464, 418–422 (2010).

7. Macho, A. P. & Zipfel, C. Plant PRRs and the activation of innate immune signaling. Mol. Cell 54, 263–272 (2014).

8. Mwimba, M. et al. Daily humidity oscillation regulates the circadian clock to influence plant physiology. Nat. Commun. 9, 4290 (2018).

9. Wang, W. et al. Timing of plant immune responses by a central circadian regulator. Nature 470, 110–114 (2011).

10. Zhou, M. et al. Redox rhythm reinforces the circadian clock to gate immune response. Nature 523, 472–476 (2015).

11. Madden, L. V. Effects of rain on splash dispersal of fungal pathogens. *CANADIAN J*. Plant Pathol. 19, 225–230 (1997).

12. Schwartz, H. F., Otto, K. L. & Gent, D. H. Relation of temperature and rainfall to development of Xanthomonas and Pantoea leaf blights of onion in Colorado. Plant Dis. 87, 11–14 (2003).

13. Melotto, M., Underwood, W., Koczan, J., Nomura, K. & He, S. Y. Plant stomata function in innate immunity against bacterial invasion. Cell 126, 969–980 (2006).

14. Xin, X. F. et al. Bacteria establish an aqueous living space in plants crucial for virulence. Nature 539, 524–529 (2016).

15. Braam, J. & Davis, R. W. Rain-, wind-, and touch-induced expression of calmodulin and calmodulin-related genes in Arabidopsis. Cell 60, 357–364 (1990).

16. Jaffe, M. J. Thigmomorphogenesis: The response of plant growth and development to mechanical stimulation : With special reference to *Bryonia dioica*. Planta 114, 143–157 (1973).

17. Sanyal, D. & Bangerth, F. Stress induced ethylene evolution and its possible relationship to auxin-transport, cytokinin levels, and flower bud induction in shoots of apple seedlings and bearing apple trees. Plant Growth Regul. 24, 124–134 (1998).

18. Johnson, K. A., Sistrunk, M. L., Polisensky, D. H. & Braam, J. *Arabidopsis thaliana* responses to mechanical stimulation do not require ETR1 or EIN2. Plant Physiol. 116, 643–649 (1998).

19. Lange, M. J. & Lange, T. Touch-induced changes in *Arabidopsis* morphology dependent on gibberellin breakdown. Nat. Plants 1, 14025 (2015).

20. Van Moerkercke, A. et al. A MYC2/MYC3/MYC4-dependent transcription factor network regulates water spray-responsive gene expression and jasmonate levels. Proc. Natl. Acad. Sci. U S A 116, 23345–23356 (2019).

21. Li, B., Meng, X., Shan, L. & He, P. Transcriptional regulation of pattern-triggered immunity in plants. Cell Host Microbe 19, 641–650 (2016).

22. Pandey, S. P. & Somssich, I. E. The role of WRKY transcription factors in plant immunity. Plant Physiol. 150, 1648–1655 (2009).

23. Dong, X., Mindrinos, M., Davis, K. R. & Ausubel, F. M. Induction of Arabidopsis defense genes by virulent and avirulent Pseudomonas syringae strains and by a cloned avirulence gene. Plant Cell 3, 61–72 (1991).

24. Fu, Z. Q. & Dong, X. Systemic acquired resistance: turning local infection into global defense. Annu. Rev. Plant Biol. 64, 839–863 (2013).

25. Howe, G. A. & Jander, G. Plant immunity to insect herbivores. Annu. Rev. Plant Biol. 59, 41–66 (2008).

26. Tena, G., Boudsocq, M. & Sheen, J. Protein kinase signaling networks in plant innate immunity. Curr. Opin. Plant Biol.. 14, 519–529 (2011).

27. Galletti, R., Ferrari, S. & De Lorenzo, G. Arabidopsis MPK3 and MPK6 play different roles in basal and oligogalacturonide- or flagellin-induced resistance against *Botrytis cinerea*. Plant Physiol. 157, 804–814 (2011).

28. Su, J. et al. Active photosynthetic inhibition mediated by MPK3/MPK6 is critical to effector-triggered immunity. PLoS Biol. 16, e2004122 (2018).

29. Bouche, N., Scharlat, A., Snedden, W., Bouchez, D. & Fromm, H. A novel family of calmodulin-binding transcription activators in multicellular organisms. J. Biol. Chem. 277, 21851–21861 (2002).

30. Doherty, C. J., Van Buskirk, H. A., Myers, S. J. & Thomashow, M. F. Roles for *Arabidopsis* CAMTA transcription factors in cold-regulated gene expression and freezing tolerance. Plant Cell 21, 972–984 (2009).

31. Finkler, A., Ashery-Padan, R. & Fromm, H. CAMTAs: calmodulin-binding transcription activators from plants to human. FEBS Lett. 581, 3893–3898 (2007).

32. Yamamoto, Y. Y. et al. Identification of plant promoter constituents by analysis of local distribution of short sequences. BMC Genomics 8, 67 (2007).

33. Yang, Y. et al. Genome-wide identification of *CAMTA* gene family members in *Medicago truncatula* and their expression during root nodule symbiosis and hormone treatments. Front. Plant Sci. 6, 459 (2015).

34. Du, L. et al. Ca(2+)/calmodulin regulates salicylic-acid-mediated plant immunity. Nature 457, 1154–1158 (2009).

35. Galon, Y. et al. Calmodulin-binding transcription activator (CAMTA) 3 mediates biotic defense responses in *Arabidopsis*. FEBS Lett. 582, 943–948 (2008).

36. Kim, Y. S. et al. CAMTA-mediated regulation of salicylic acid immunity pathway genes in Arabidopsis exposed to low temperature and pathogen infection. Plant Cell 29, 2465–2477 (2017).

37. Nie, H. et al. SR1, a calmodulin-binding transcription factor, modulates plant defense and ethylene-induced senescence by directly regulating NDR1 and EIN3. Plant Physiol. 158, 1847–1859 (2012).

38. Kim, Y., Park, S., Gilmour, S. J. & Thomashow, M. F. Roles of CAMTA transcription factors and salicylic acid in configuring the low-temperature transcriptome and freezing tolerance of Arabidopsis. Plant J. 75, 364–376 (2013).

39. Thorvaldsdottir, H., Robinson, J. T. & Mesirov, J. P. Integrative Genomics Viewer (IGV): high-performance genomics data visualization and exploration. Brief Bioinformatics 14, 178–192 (2013).

40. Gutschick, V. P. Biotic and abiotic consequences of differences in leaf structure. New Phytol. 143, 3–18 (2002).

41. Wagner, G. J., Wang, E. & Shepherd, R. W. New approaches for studying and exploiting an old protuberance, the plant trichome. Ann. Bot. 93, 3–11 (2004).

42. Zhou, L. H. et al. The Arabidopsis trichome is an active mechanosensory switch. Plant Cell Environ. 40, 611–621 (2017).

43. Tian, L. et al. Imaging neural activity in worms, flies and mice with improved GCaMP calcium indicators. Nat. Methods 6, 875–881 (2009).

44. Toyota, M. et al. Glutamate triggers long-distance, calcium-based plant defense signaling. Science 361, 1112–1115 (2018).

45. Larkin, J. C., Oppenheimer, D. G., Lloyd, A. M., Paparozzi, E. T. & Marks, M. D. Roles of the GLABROUS1 and TRANSPARENT TESTA GLABRA genes in Arabidopsis trichome development. Plant Cell 6, 1065–1076 (1994).

46. Xia, Y. et al. The *glabra1* mutation affects cuticle formation and plant responses to microbes. Plant Physiol. 154, 833–846 (2010).

47. Mao, G. et al. Phosphorylation of a WRKY transcription factor by two pathogen-responsive MAPKs drives phytoalexin biosynthesis in *Arabidopsis*. Plant Cell 23, 1639–1653 (2011).

48. Tsuda, K. & Somssich, I. E. Transcriptional networks in plant immunity. New Phytol. 206, 932–947 (2015).

49. Chehab, E. W., Yao, C., Henderson, Z., Kim, S. & Braam, J. *Arabidopsis* touch-induced morphogenesis is jasmonate mediated and protects against pests. Curr. Biol. 22, 701–706 (2012).

50. Bauer, H. et al. The contribution of bacteria and fungal spores to the organic carbon content of cloud water, precipitation and aerosols. Atmospheric Research 64, 109–119 (2002).

51. Prussin, A. J. 2nd, Marr, L. C. & Bibby, K. J. Challenges of studying viral aerosol metagenomics and communities in comparison with bacterial and fungal aerosols. FEMS Microbiol. Lett. 357, 1–9 (2014).

52. Huynh, T.-P. & Haick, H. Learning from an intelligent mechanosensing system of plants. *Adv*. Mater. Technol. 4, 1800464 (2019).

53. Scherzer, S., Federle, W., Al-Rasheid, K. A. S. & Hedrich, R. Venus flytrap trigger hairs are micronewton mechano-sensors that can detect small insect prey. Nat. Plants 5, 670–675 (2019).

54. Bacete, L. & Hamann, T. The Role of mechanoperception in plant cell wall integrity maintenance. Plants (Basel, Switzerland) 9, (2020).

55. Suda, H. et al. Calcium dynamics during trap closure visualized in transgenic Venus flytrap. Nat. Plants 6, 1219–1224 (2020).

56. Betsuyaku, S. et al. Salicylic acid and jasmonic acid pathways are activated in spatially different domains around the infection site during effector-triggered immunity in *Arabidopsis thaliana*. Plant Cell Physiol. 59, 8–16 (2018).

57. Murashige, T. & Skoog, F. A revised medium for rapid growth and bio assays with Tobacco tissue cultures. Physiol. Plant. 15, 473–497 (1962).

58. Gunn, R. & Kinzer, G. D. The terminal velocity of fall for water droplet in stagnant air. J. Atmospheric Sci. 6, 243–248 (1949).

59. Robinson, M. D., McCarthy, D. J. & Smyth, G. K. edgeR: a Bioconductor package for differential expression analysis of digital gene expression data. Bioinformatics 26, 139–140 (2010).

60. Langmead, B. & Salzberg, S. L. Fast gapped-read alignment with Bowtie 2. Nat. Methods 9, 357–359 (2012).

61. Maere, S., Heymans, K. & Kuiper, M. BiNGO: a Cytoscape plugin to assess overrepresentation of gene ontology categories in biological networks. Bioinformatics 21, 3448–3449 (2005).

62. Denoux, C. et al. Activation of defense response pathways by OGs and flg22 elicitors in *Arabidopsis* seedlings. Mol. Plant 1, 423–445 (2008).

63. Matsuura, T., Mori, I. C., Himi, E. & Hirayama, T. Plant hormone profiling in developing seeds of common wheat (*Triticum aestivum L*.). Breeding Science 69, 601–610 (2019).

64. Yamaguchi, N. et al. PROTOCOLS: Chromatin immunoprecipitation from Arabidopsis tissues. The arabidopsis book 12, e0170 (2014).

65. Li, H. et al. The Sequence Alignment/Map format and SAMtools. Bioinformatics 25, 2078–2079 (2009).

66. Zhang, Y. et al. Model-based Analysis of ChIP-Seq (MACS). Genome Biol. 9, R137 (2008).

67. Quinlan, A. R. & Hall, I. M. BEDTools: a flexible suite of utilities for comparing genomic features. Bioinformatics 26, 841–842 (2010).

68. Machanick, P. & Bailey, T. L. MEME-ChIP: motif analysis of large DNA datasets. Bioinformatics 27, 1696–1697 (2011).

69. Cao, H., Bowling, S. A., Gordon, A. S. & Dong, X. Characterization of an Arabidopsis mutant that is nonresponsive to inducers of systemic acquired resistance. Plant Cell 6, 1583–1592 (1994).

70. Hiruma, K. et al. *Arabidopsis ENHANCED DISEASE RESISTANCE 1* is required for pathogen-induced expression of plant defensins in nonhost resistance, and acts through interference of *MYC2*-mediated repressor function. Plant J. 67, 980–992 (2011).

