## Supplementary_Figures for "Mechanosensory trichome cells evoke a mechanical stimuli–induced immune response in plants"

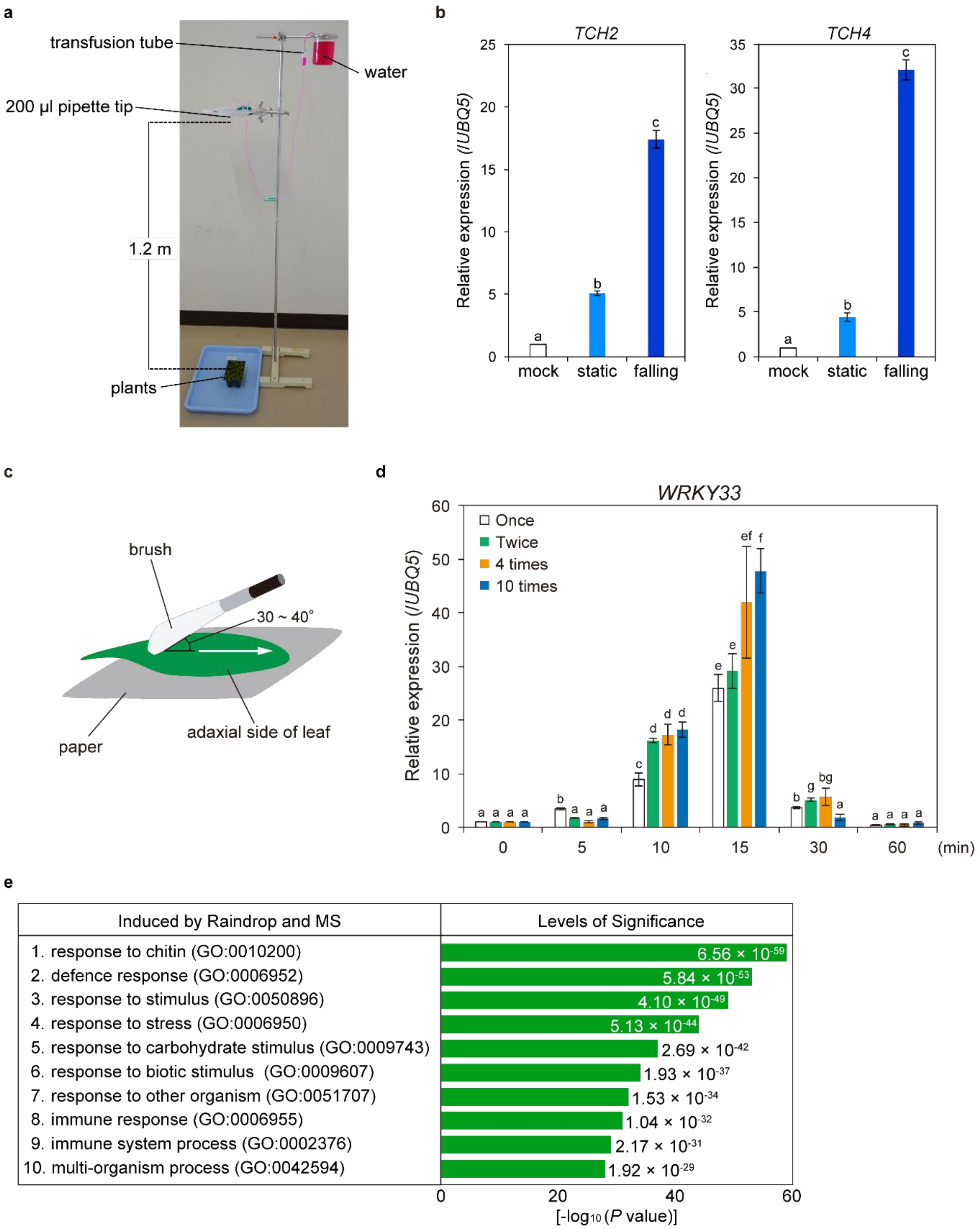
a
transfusion tube
water
200 µl pipette tip
1.2 m
plants
b
TCH2
TCH4
Relative expression (/UBQ5)
mock
static
falling
c
brush
30 ~ 40°
paper
adaxial side of leaf
d
WRKY33
Once
Twice
4 times
10 times
Relative expression (/UBQ5)
(min)
e
Induced by Raindrop and MS
Levels of Significance
1. response to chitin (GO:0010200)
2. defence response (GO:0006952)
3. response to stimulus (GO:0050896)
4. response to stress (GO:0006950)
5. response to carbohydrate stimulus (GO:0009743)
6. response to biotic stimulus (GO:0009607)
7. response to other organism (GO:0051707)
8. immune response (GO:0006955)
9. immune system process (GO:0002376)
10. multi-organism process (GO:0042594)
6.56 × 10-59
5.84 × 10-53
4.10 × 10-49
5.13 × 10-44
2.69 × 10-42
1.93 × 10-37
1.53 × 10-34
1.04 × 10-32
2.17 × 10-31
1.92 × 10-29
[-log10 (P value)]

**Supplementary Fig. 1 | Raindrop-induced MS initiate the transient expression of defence-related genes. a** The device drops 13 μL water droplets from a height of 1.2 m onto the adaxial side of leaves. **b** The adaxial side of wild-type leaves was treated with water droplets and collected after 15 min. The expression levels of *TCH2* and *TCH4* were significantly higher in response to one falling raindrop (falling) than to one water droplet directly placed on the leaf surface (static). Data are presented as mean ± SD. Different letters above bars indicate significant differences ($P < 0.05$). **c** Brushing method. The adaxial side of leaves from 4-week-old plants was brushed four times along the main vein at an angle of 30-40°. **d** The adaxial side of wild-type leaves was brushed for the indicated number of times. The maximum level of *WRKY33* expression was detected 15 min after brushing. Mean ± SD. Different letters above bars indicate significant differences ($P < 0.05$). **e** Gene Ontology (GO) categories enriched in 917 genes induced by brushing. Enrichment of GO categories for the biological process was determined using BiNGO. The table shows GO terms with $P$ values from the lowest to the tenth.

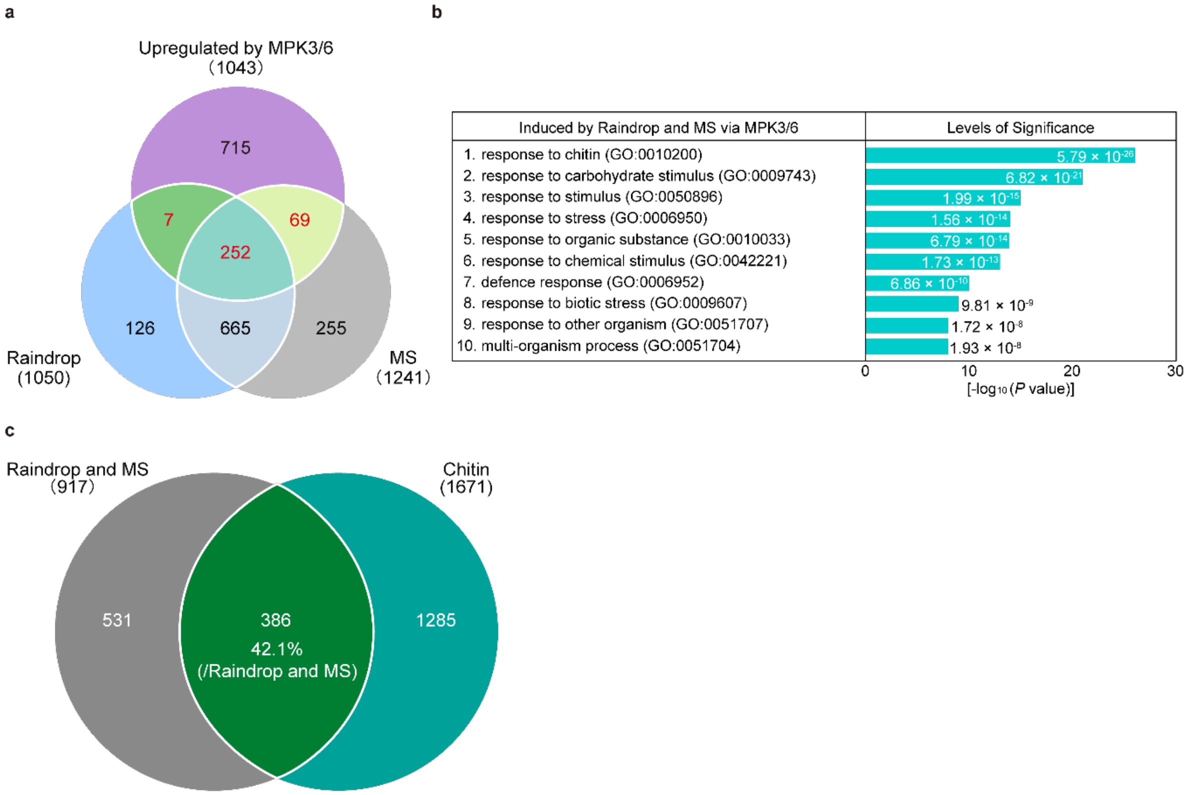


**Supplementary Fig. 2 | MS induces the expression of defence-related genes through the activation of MPK3/MPK6. a** Venn diagram of the overlap between genes upregulated by raindrops (1,050 genes), MS (1,241 genes), and MPK3/MPK6 (1,043 genes). A total of 328 genes (shown in red) are present in at least three groups ($P < 0.05$). **b** Enriched GO categories of 328 genes induced by raindrops and MS through the activation of MPK3/MPK6 shown in (**a**). The top 10 categories are shown in ascending order of *P* values. **c** Venn diagram of the overlap between transcriptome datasets from raindrop- and MS-induced genes (917 genes) and chitin-induced genes (1,671 genes) ($P < 0.05$).

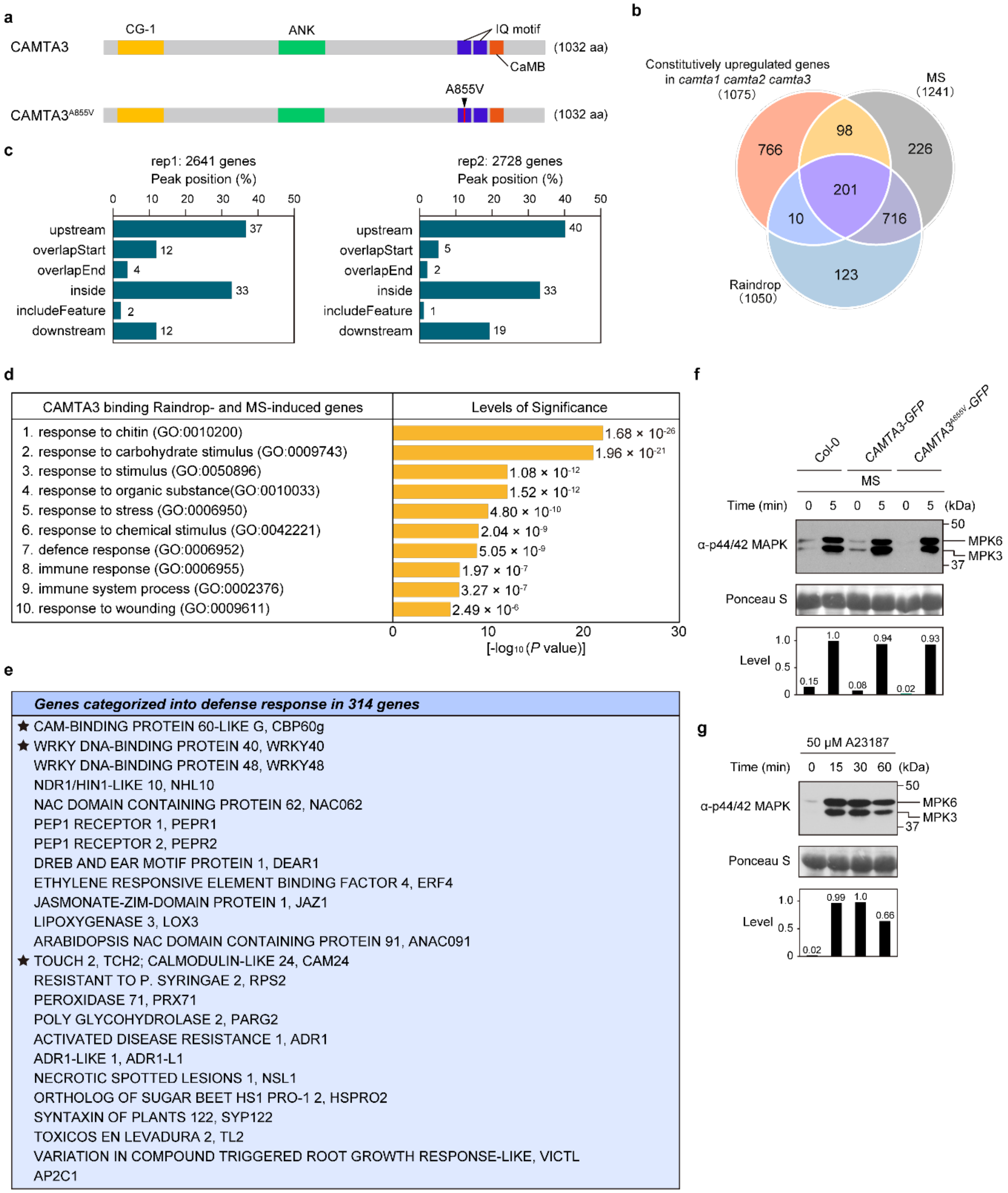
a
CAMTA3
CG-1
ANK
IQ motif
CaMB
(1032 aa)
A855V
CAMTA3A855V
(1032 aa)
b
Constitutively upregulated genes in camta1 camta2 camta3 (1075)
MS (1241)
Raindrop (1050)
766
98
226
201
10
716
123
c
rep1: 2641 genes
Peak position (%)
upstream 37
overlapStart 12
overlapEnd 4
inside 33
includeFeature 2
downstream 12
rep2: 2728 genes
Peak position (%)
upstream 40
overlapStart 5
overlapEnd 2
inside 33
includeFeature 1
downstream 19
d
CAMTA3 binding Raindrop- and MS-induced genes
Levels of Significance
1. response to chitin (GO:0010200) 1.68 × 10-26
2. response to carbohydrate stimulus (GO:0009743) 1.96 × 10-21
3. response to stimulus (GO:0050896) 1.08 × 10-12
4. response to organic substance(GO:0010033) 1.52 × 10-12
5. response to stress (GO:0006950) 4.80 × 10-10
6. response to chemical stimulus (GO:0042221) 2.04 × 10-9
7. defence response (GO:0006952) 5.05 × 10-9
8. immune response (GO:0006955) 1.97 × 10-7
9. immune system process (GO:0002376) 3.27 × 10-7
10. response to wounding (GO:0009611) 2.49 × 10-6
[-log10 (P value)]
e
Genes categorized into defense response in 314 genes
★ CAM-BINDING PROTEIN 60-LIKE G, CBP60g
★ WRKY DNA-BINDING PROTEIN 40, WRKY40
WRKY DNA-BINDING PROTEIN 48, WRKY48
NDR1/HIN1-LIKE 10, NHL10
NAC DOMAIN CONTAINING PROTEIN 62, NAC062
PEP1 RECEPTOR 1, PEPR1
PEP1 RECEPTOR 2, PEPR2
DREB AND EAR MOTIF PROTEIN 1, DEAR1
ETHYLENE RESPONSIVE ELEMENT BINDING FACTOR 4, ERF4
JASMONATE-ZIM-DOMAIN PROTEIN 1, JAZ1
LIPOXYGENASE 3, LOX3
ARABIDOPSIS NAC DOMAIN CONTAINING PROTEIN 91, ANAC091
★ TOUCH 2, TCH2; CALMODULIN-LIKE 24, CAM24
RESISTANT TO P. SYRINGAE 2, RPS2
PEROXIDASE 71, PRX71
POLY GLYCOHYDROLASE 2, PARG2
ACTIVATED DISEASE RESISTANCE 1, ADR1
ADR1-LIKE 1, ADR1-L1
NECROTIC SPOTTED LESIONS 1, NSL1
ORTHOLOG OF SUGAR BEET HS1 PRO-1 2, HSPRO2
SYNTAXIN OF PLANTS 122, SYP122
TOXICOS EN LEVADURA 2, TL2
VARIATION IN COMPOUND TRIGGERED ROOT GROWTH RESPONSE-LIKE, VICTL
AP2C1
f
Col-0
CAMTA3-GFP
CAMTA3A855V-GFP
MS
Time (min)
(kDa)
α-p44/42 MAPK
MPK6
MPK3
Ponceau S
Level
0.15
1.0
0.08
0.94
0.02
0.93
g
50 μM A23187
Time (min)
(kDa)
α-p44/42 MAPK
MPK6
MPK3
Ponceau S
Level
0.02
0.99
1.0
0.66

**Supplementary Fig. 3 | MS-induced genes regulated by CAMTAs show enrichment in categories associated with plant immunity. a** Diagram of CAMTA3 and CAMTA3$^{A855V}$ proteins. CG-1, DNA (CGCG box)-binding domain; ANK, ankyrin repeat domain; IQ and CaMB, calmodulin (CaM)-binding domain. **b** Venn diagram of transcriptome datasets obtained by brushing (1,241 genes) and raindrops (1,050 genes) and constitutively upregulated genes in the *camta1 camta2 camta3* triple mutant (1,075 genes) ($P < 0.05$). Overlap with MS-induced genes; constitutively upregulated genes in the *camta1 camta2 camta3* triple mutant (28.7%; 309/1,075 genes). The 201 shared upregulated genes include *WRKY33*, *WRKY40*, and *CBP60g*. **c** Positions of the 2,641 and 2,728 CAMTA3-binding peaks in 2,011 genes relative to the annotated nearest transcription start site (TSS). Upstream, peak resides upstream of the feature; overlapStart, peak overlaps with the start of the feature; overlapEnd, peak overlaps with the end of the feature; inside, peak resides inside the feature; includeFeature, peak includes the feature entirely; downstream, peak resides downstream of the feature. **d** Enriched Gene Ontology categories of 314 CAMTA3 target genes shown in Fig. 3d. The top 10 categories are shown in ascending order of *P* values. **e** Representative MS-induced defence genes in 314 CAMTA target genes. Black stars indicate the RT-qPCR marker genes used in this work. **f** Brush treatment (MS) induces MAPK activation in Col-0, *camta2 camta3 CAMTA3-GFP*, and *camta2 camta3 CAMTA3*$^{A855V}$*-GFP*. Total proteins were extracted from 4-week-old plants 5 min after MS treatment (4 brushing) and detected by immunoblot analysis with anti-p44/42 MAPK antibodies. Relative phosphorylation levels are shown below each blot. **g** Cytosolic $Ca^{2+}$-induced MAPK activation in Col-0. Total proteins were extracted from 12-day-old plants treated with 50 µM calcium ionophore A23187 and detected by immunoblot analysis with anti-p44/42 MAPK antibodies. Relative phosphorylation levels are shown below each blot.

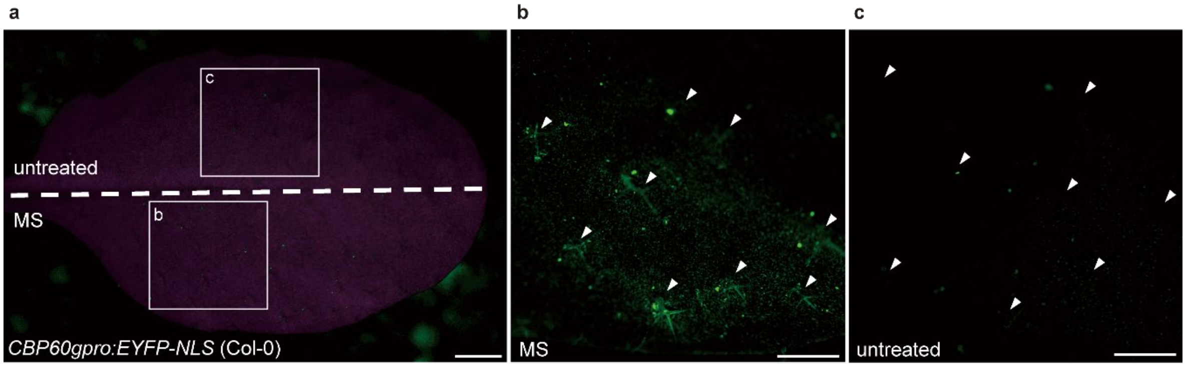


**Supplementary Fig. 4 | MS induces the expression of *CBP60g* in cells surrounding trichomes. a-c** YFP fluorescence from the whole leaf of *CBP60gpro:EYFP-NLS* (Col-0) with (MS; bottom half) or without brushing (untreated; top half) (**a**), with zoomed-in views of MS (**b**) and untreated (**c**) areas. Arrowheads indicate trichomes (**b**, **c**). Scale bars, 0.5 mm (**a**), 0.3 mm (**b**, **c**).

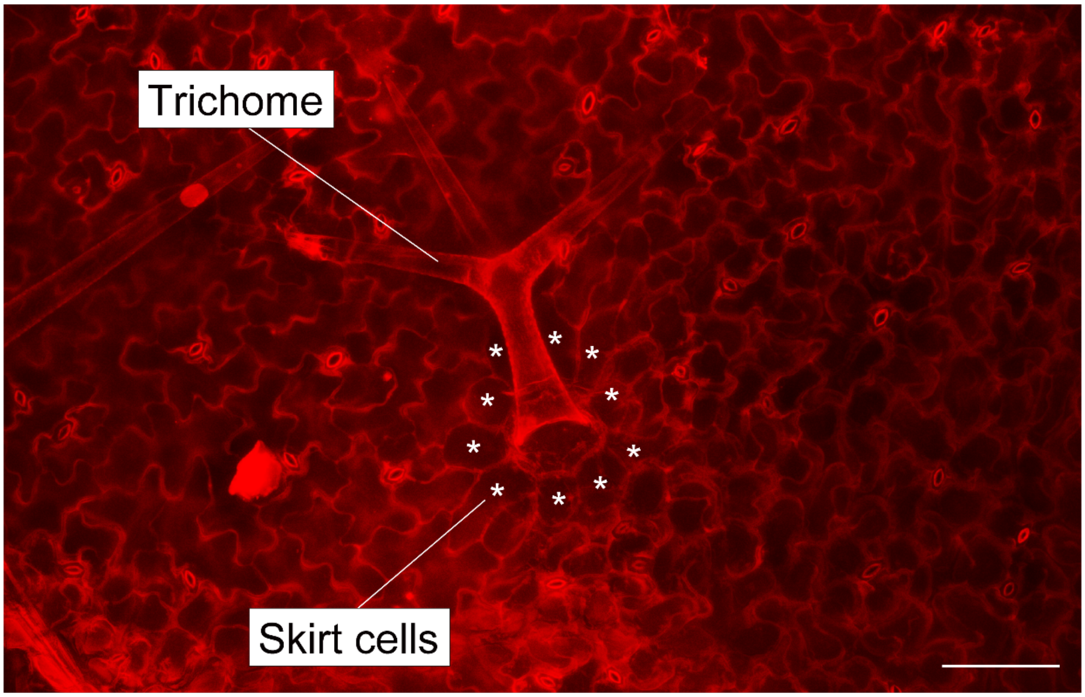


**Supplementary Fig. 5 | Trichomes are surrounded by their skirt cells.** Confocal imaging showing propidium iodide staining of the leaf cell wall. Asterisks indicate skirt cells. Trichome is surrounded by skirt cells, and these cells are surrounded by epidermal cells. Scale bars, 0.1 mm.

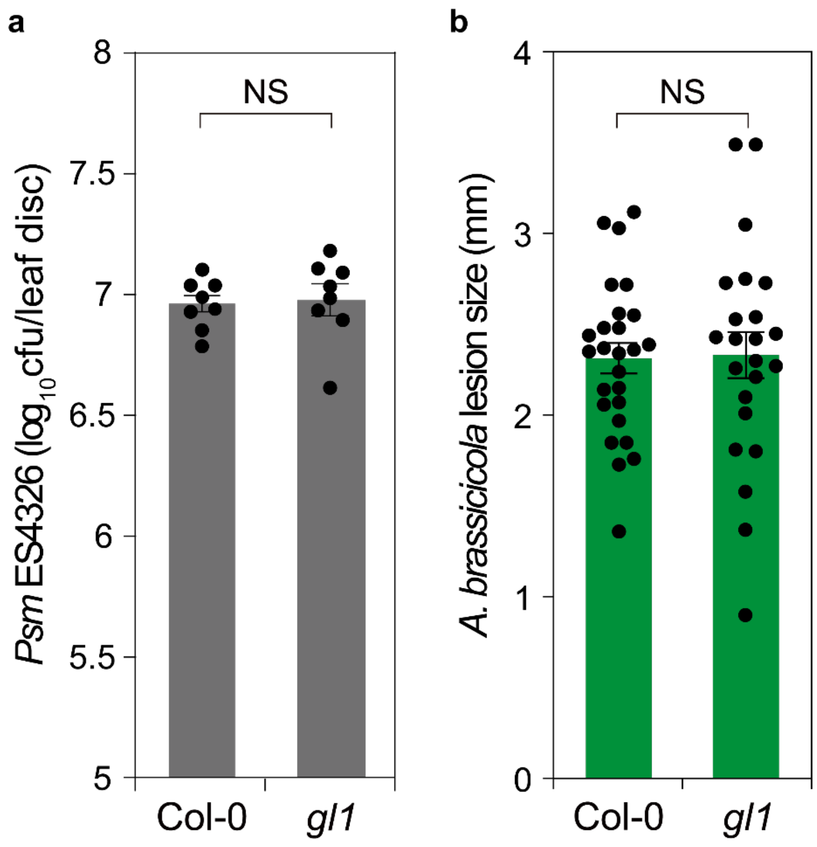


**Supplementary Fig. 6 | The local resistance against *Psm* ES4326 and *Alternaria brassicicola* Ryo-1 are normally induced in our mock condition. a** Growth of *Psm* ES4326 in Col-0 and *gl1* leaves 2 days after inoculation. Error bars represent SE. NS, not significant. Cfu, colony-forming units. **b** Disease progression of *A. brassicicola* in Col-0 and *gl1* leaves 3 days after inoculation. Error bars represent SE. NS, not significant. Scale bar, 0.5 mm.

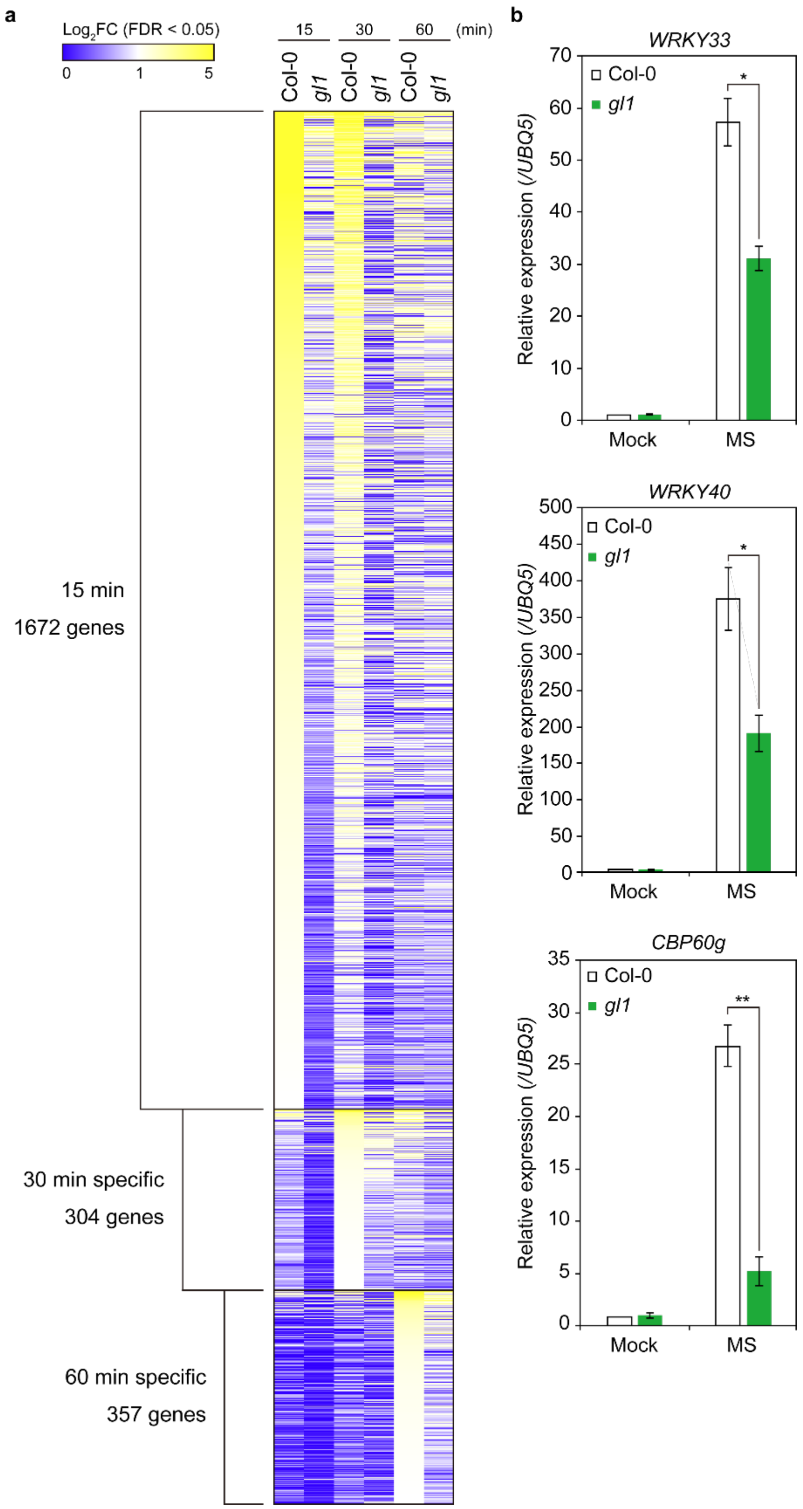
a
Log2FC (FDR < 0.05)
0
1
5
15
30
60
(min)
Col-0
gl1
Col-0
gl1
Col-0
gl1
15 min
1672 genes
30 min specific
304 genes
60 min specific
357 genes
b
WRKY33
Relative expression (/UBQ5)
Col-0
gl1
Mock
MS
WRKY40
Relative expression (/UBQ5)
Col-0
gl1
Mock
MS
CBP60g
Relative expression (/UBQ5)
Col-0
gl1
Mock
MS

**Supplementary Fig. 7 | Trichomes amplify the expression of MS-induced genes. a** Heatmap of differentially expressed genes obtained from transcriptome datasets for Col-0 and *gl1* plants treated by MS (4 brushing). Genes upregulated 15 min after brushing: 1,672 genes; unique induced 30 min after brushing: 304 genes; unique induced 60 min after brushing: 357 genes. See also Fig. 5c, d. **b** Four-week-old Col-0 and *gl1* plants were brushed four times. Transcript levels of *WRKY33*, *WRKY40*, and *CBP60g* 15 min after brushing were determined using RT-qPCR and normalized to *UBQ5*. Data are presented as mean ± SD. Asterisks indicate significant difference (one- and two-way ANOVA; *$P < 0.05$, **$P < 0.01$).

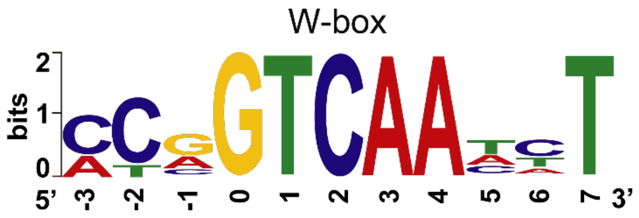


**Supplementary Fig. 8 | Promoter analysis of MAPK-regulated 252 genes among 917 raindrop- and MS-induced genes.** The WRKY-binding W-box (TTGACC) was overrepresented among these genes.
